# Structural basis of the catalytic and allosteric mechanism of bacterial acetyltransferase PatZ

**DOI:** 10.1101/2024.11.12.623305

**Authors:** Jun Bae Park, Gwanwoo Lee, Yu-Yeon Han, Dongwook Kim, Kyoo Heo, Jeesoo Kim, Juhee Park, Hyosuk Yun, Chul Won Lee, Hyun-Soo Cho, Jong-Seo Kim, Martin Steinegger, Yeong-Jae Seok, Soung-Hun Roh

**Affiliations:** School of Biological Sciences, Seoul National University, Seoul, 08826, Republic of Korea; Institute of Molecular Biology and Genetics, Seoul National University, Seoul, 08826, Republic of Korea; Institute of Microbiology, Seoul National University, Seoul, 08826, Republic of Korea; Interdisciplinary Program in Bioinformatics, Seoul National University, Seoul, 08826, Republic of Korea; Center for RNA Research, Institute for Basic Science, Seoul 08826, Republic of Korea; Department of Chemistry, Chonnam National University, Gwangju, 61186, Republic of Korea; Department of Systems Biology, College of Life Science and Biotechnology, Yonsei University, 50 Yonsei-ro, Seoul, 03722, Republic of Korea; Artificial Intelligence Institute, Seoul National University, Seoul, 08826, Republic of Korea

**Keywords:** PatZ, YfiQ, Acetyltransferase, Allosterism, CryoEM

## Abstract

GCN5-related *N*-Acetyltransferases (GNATs) play a crucial role in regulating bacterial metabolism by acetylating specific target proteins. Despite their importance in bacterial physiology, the mechanisms underlying GNATs’ enzymatic and regulatory functions remain poorly understood. In this study, we elucidated the structures of *Escherichia coli* PatZ, a type I GNAT, and investigated its ligand interactions, catalytic processes, and allosterism. PatZ functions as a homotetramer, with each subunit comprising a catalytic domain and a regulatory domain. Our findings reveal that the regulatory domain is essential for acetyltransferase activity, as it not only induces cooperative conformational changes in the catalytic domain but also directly contributes to the formation of substrate binding pockets. Furthermore, a protein structure-based analysis on the evolution of bacterial GNAT types reveals a distinct pattern of the regulatory domain across phyla, underscoring the regulatory domain’s critical role in responding to cellular energy status.

**SIGNIFICANCE STATEMENT:** Post-translational modifications, particularly acetylation mediated by GCN5-related N-Acetyltransferases (GNATs), play a crucial role in bacterial physiology. Protein acetyltransferase Z (PatZ) is a key GNAT with diverse substrates, essential for understanding the bacterial acetylome. This study employs cryogenic electron microscopy, X-ray crystallography, and biochemical analyses to elucidate the mechanistic regulation of *Escherichia coli* PatZ. Our high-resolution structures reveal PatZ’s homo-tetrameric architecture, with each subunit comprising regulatory and GNAT domains. We characterize ligand-PatZ interactions, demonstrating ligand-induced conformational changes that facilitate allosteric regulation of the catalytic domain. Furthermore, our analyses elucidate the regulatory domain’s contribution to substrate binding pocket formation, potentially enhancing substrate specificity. Structure-based phylogenetic analysis provides insights into the evolution of diverse regulatory domains in the GNAT superfamily across bacterial taxonomy. This first visualization of PatZ advances our mechanistic understanding of bacterial physiology, offering novel insights into GNAT-mediated bacterial adaptations.

**HIGHLIGHTS:**

- *E. coli* PatZ forms a homotetramer, with each subunit possessing a GNAT catalytic domain and a regulatory domain.
- Cooperative binding of acetyl-CoA to the regulatory domains is a prerequisite for inducing the structural compatibility of the catalytic domain with a substrate.
- Diverse regulatory domains in GNATs evolved to adapt to varied metabolic conditions across bacterial taxonomy.

## INTRODUCTION

Post-translational modifications (PTMs) play a crucial role in modifying protein structure, stability, activity, and their placement within the cell^1–3^. Acetylation stands out as one of the predominant PTMs, involving the attachment of an acetyl group to the lysine residues or the protein’s N-terminal^4^. This modification is a widespread phenomenon across all forms of life, facilitated either enzymatically by acetyltransferases or non-enzymatically via high-energy compounds such as acetyl phosphate (AcP) or acetyl-CoA (AcCoA)^5–8^. In eukaryotes, acetylation affects both histones and non-histone proteins^9,10^, thereby influencing a broad array of cellular and physiological processes including transcription, phase separation, autophagy, mitosis, cellular differentiation, and neuronal activities^11–16^ and disruption in acetylation regulation is linked to cancer development and aging^17,18^. In prokaryotes, acetylation plays a significant role in regulating metabolism, modifying metabolic fluxes, cell growth, and survival^19–21^. It also plays a part in the regulation of gene expression by altering the activity of transcription factors according to environmental shifts^22^. While the physiological significance of *N*ε-lysine acetylation is widely recognized in eukaryotes, its biological relevance in bacteria is still emerging.

Enzymatic acetylation involves the addition of an acetyl group to lysine residues by enzymes known as lysine acetyltransferases (KATs)^23^. Based on sequence homology and biochemical characteristics, KATs are classified into several families, including the p300/cAMP response element binding protein binding protein (CBP) family, Moz, Ybf2/Sas3, Sas2, Tip60 (MYST) family, and GCN5-related N-acetyltransferase (GNAT) family^11,24,25^. The MYST and p300/CBP families are found exclusively in eukaryotic cells, whereas the GNAT family includes orthologous proteins present in bacteria, eukaryotes, and archaea^23^. Approximately 65% of the GNAT domain superfamily is found in bacteria^26^. Recent studies reveal that acetylation significantly impacts bacterial physiology, influencing translation in *Escherichia coli*^27^, metabolic flux in *Salmonella enterica*^28^, and virulence in *Francisella novicida*^29^. These findings underscore the broad and vital impact of acetylation on bacterial physiology, ranging from basic cellular processes to complex pathogenic mechanisms.

The catalytic GNAT domain is highly conserved across species. However, bacterial GNATs exhibit unique regulatory domains which are diverse in different bacterial species^8^. These GNATs are classified into five types based on their architecture and arrangement of catalytic GNAT domain and regulatory domains (Figure 1a). Type I and II feature a homologous structure of NDP-forming acyl-CoA synthetase in the regulatory domain, while Type III has a cAMP-binding, ACT (aspartate kinase, chorismate mutase, TyrA) or NADP^+^-binding protein^8^. Types IV and V do not possess regulatory domain. Truncation or mutations in these regulatory domains significantly diminished GNAT-mediated catalytic activity^8,30^, implying that the regulatory and GNAT domains are functionally interdependent. In particular, protein acetyltransferase Z (PatZ), also known as YfiQ, is one of the most well characterized Type I GNATs, with approximately 80% of its amino acids comprising the regulatory domain, and 20 % GNAT domain. PatZ alters its oligomeric states and activity based on the presence of AcCoA, showing different patterns in *E. coli* and *S. enterica*^31,32^. Recent studies about PatZ demonstrate that regulatory domain increases the acetyltransferase activity and that ligand binding to regulatory domain elevates AcCoA affinity to GNAT domain^31^. However, the precise mechanisms remain unclear.

**Figure 1.**
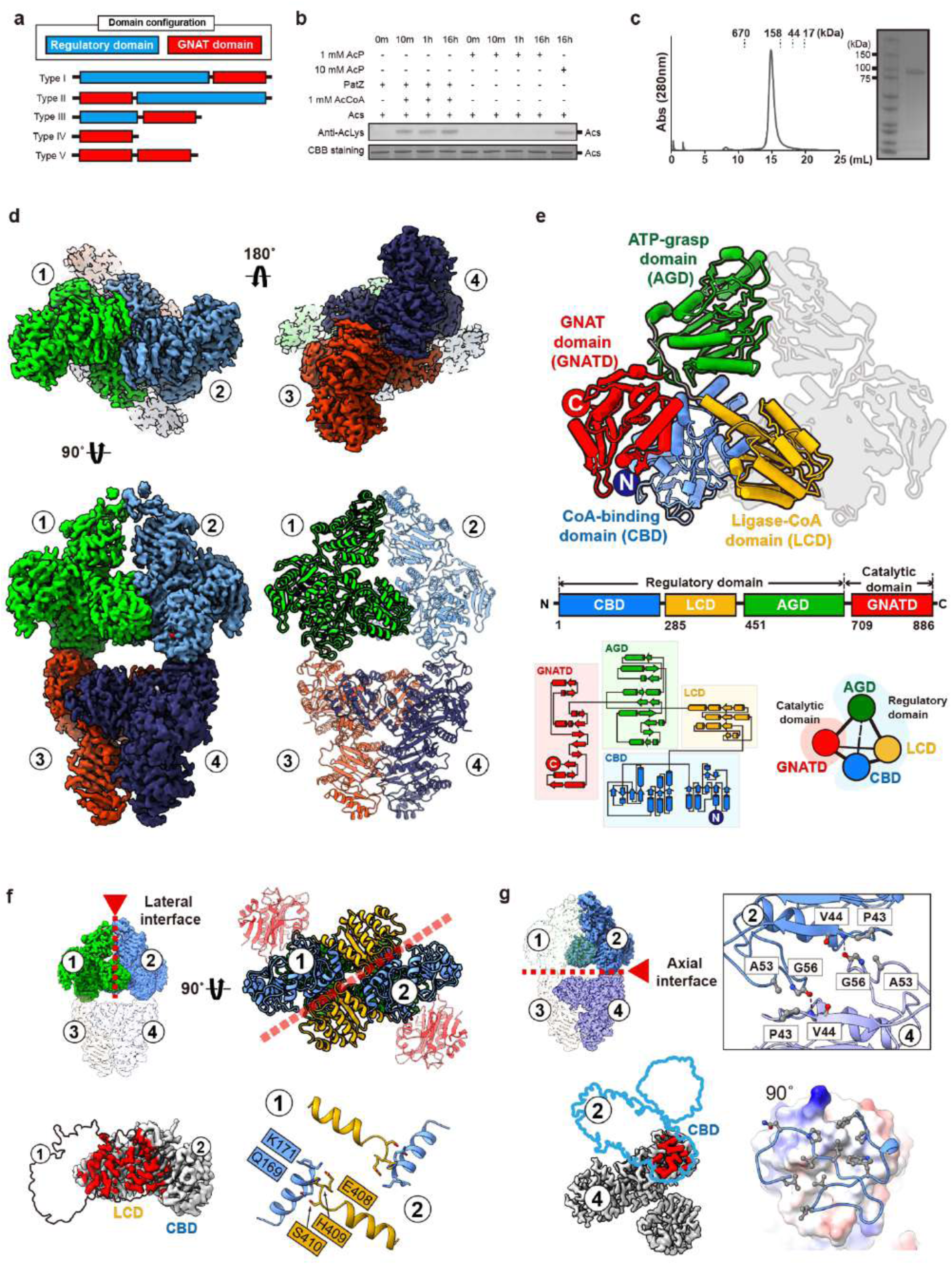
Structure of the apo-PatZ. **a,** Domain architecture of bacterial Nε-lysine acetyltransferase that includes the GNAT domain. **b,** Comparison of enzymatic acetylation by PatZ and nonenzymatic acetylation by acetyl-phosphate. Western blot analysis using anti-acetylated lysine antibody. CBB denotes coomassie brilliant blue. **c,** Analysis of the tetrameric state and purity of PatZ using size exclusion chromatography and SDS-PAGE. **d,** CryoEM map and refined model. Each color denotes a distinct subunit. **e,** The structural arrangement of PatZ, consisting of four subdomains**. f,** The interaction between subunits or subdomains forming the lateral interface **g,** and axial interface.

In this study, we elucidated the homotetrameric structure of functional PatZ using cryo-electron microscopy (cryoEM). Each subunit of PatZ comprises four structural domains, binding to two AcCoA molecules, one ATP, and one phosphate. The unliganded-PatZ adopts an inactive conformation characterized by a closed substrate binding pocket. Our findings reveal that the regulatory domains undergo conformational changes upon ligand binding, altering their interactions with the GNAT domain. Specifically, the binding of AcCoA to the regulatory domain triggers a cooperative structural rearrangement in the GNAT domain, which facilitates the binding of the AcCoA donor and organizes the substrate in close proximity for catalysis. Furthermore, our analysis suggests that PatZ coevolved with its substrate, with the regulatory domain playing a crucial role in substrate interaction. Additionally, we conducted a phylogenetic analysis of bacterial GNAT-mediated acetyltransferase families, which provides insights into the regulatory domain’s role in activating distinct acetyl-PTM responses to various metabolic signatures.

## RESULTS

### Overall architecture of *E. coli* PatZ

To examine the biochemical and structural characteristics of PatZ, we purified the *E. coli* PatZ protein and analyzed its acetyltransferase activity towards purified *E. coli* AcCoA synthetase (Acs) using western blotting with an acetyl-lysine-specific antibody (Figure 1b). When supplied with AcCoA, PatZ efficiently catalyzed the acetylation of Acs, consistent with findings from previous studies^33,34^. This PatZ-mediated enzymatic reaction is also significantly faster than the AcP-mediated non-enzymatic reaction (Figure 1b)^35^, confirming that the prepared PatZ is active and functional.

We then delved into the three-dimensional structure of the unliganded form of PatZ to delineate its molecular architecture and assembly. The protein, with a protomer mass of approximately 100 KDa, predominantly forms a homotetramer in solution, as inferred by size exclusion chromatography (Figure 1c). CryoEM imaging of the eluted PatZ protein revealed a tetrameric structure characterized by a diamond-shaped, four-fold symmetric architecture. We then achieved a resolution of 2.52 Å for a consensus map showing the composition of four distinct subunits. These units exhibit variable resolution across the structure’s periphery and thus structural features were enhanced by multiple rounds of 3D-classification and variability analysis for each domain (Extended Data Figure 1a-e, Supplementary Table 1).

Molecular model was built by chasing the side densities and aligns closely with the cryoEM map, as demonstrated by high-quality statistical validation (Q-score=0.67, Extended Data Figure 1f). Four subunits are positioned in a cross-like arrangement, with a 180-degree rotation relative to adjacent units (Figure 1d, Supplementary Video 1), giving the tetramer lengths of approximately 190 Å and 130 Å along the long and short axes, respectively. Notably each subunit is composed of four distinct structural domains. Reflecting on similarities with NDP-forming AcCoA synthetases (ACDs)^36^ and proteins containing GNAT motifs^37^, we designated these regions accordingly the CoA binding domain (CBD), ligase-CoA domain (LCD), ATP grasp domain (AGD), and GNAT domain (GNATD), sequentially arranged from the N- to the C-terminus (Figure 1e, Supplementary Video 1).

PatZ presents a propensity to form a stable homotetramer structure, showcasing a sophisticated interaction among four protomers. The interactions exhibited significant contact areas with considerable stabilization energies, notably 3103.4 Å^2^ with a ΔiG of -28.4 kcal/mol for the lateral interface and 619.5 Å^2^ with a ΔiG of -12.0 kcal/mol for the axial interface. The formation of the lateral interface is achieved through an antiparallel hetero-domain interaction between the CBD and LCD via dominant polar interactions (Figure 1f). Conversely, the axial interface with the CBDs stacking in a back-to-back fashion at a 60-degree angle (Figure 1g). This mode of interaction enhances the stability through symmetric predominant hydrophobic contacts. An attempt to disrupt tetramer form by mutating residues at the axial interface led to significant protein aggregation. Collectively, our investigation into the PatZ protein delineates its acetyltransferase function within a uniquely stable tetrameric architecture.

### Structural basis of ligand binding to PatZ

Based on the structural homology with ACD1^36^ and GNAT^37^ proteins, we have identified four hypothetical ligand binding pockets for two AcCoAs, ATP and phosphate (or AcP). To characterize the relationship between ligand and PatZ, we incubated purified PatZ with ATP, AcCoA, and phosphate, and then analyzed it using cryoEM. We obtained a significantly higher resolution map converging at 1.99 Å (Q score=0.77, Extended Data Figure 2a-f, Supplementary Table 1). The overall shape and subunit arrangement of liganded PatZ remain identical but notably, we identified two AcCoA in each protomer, which is one in the N-terminal regulatory domain and the other in GNATD (Figure 2a,b, Supplementary Video 2). We also identified phosphate densities located at each four lateral interfaces between the LCD and CBD derived from different protomers (Figure 2a,b). Of note, we could not visualize ATP in AGD of the cryoEM map, which is potentially reasoned by structural dynamics in the domain. Therefore, we performed complementary X-ray crystallographic experiments for the AGD and confirmed bound ATP in AGD resolution with 2.24 Å (Figure 2a,b, Extended Data Figure 2g, Supplementary Table 2). Collectively, we successfully visualized all possible ligand binding structures.

**Figure 2.**
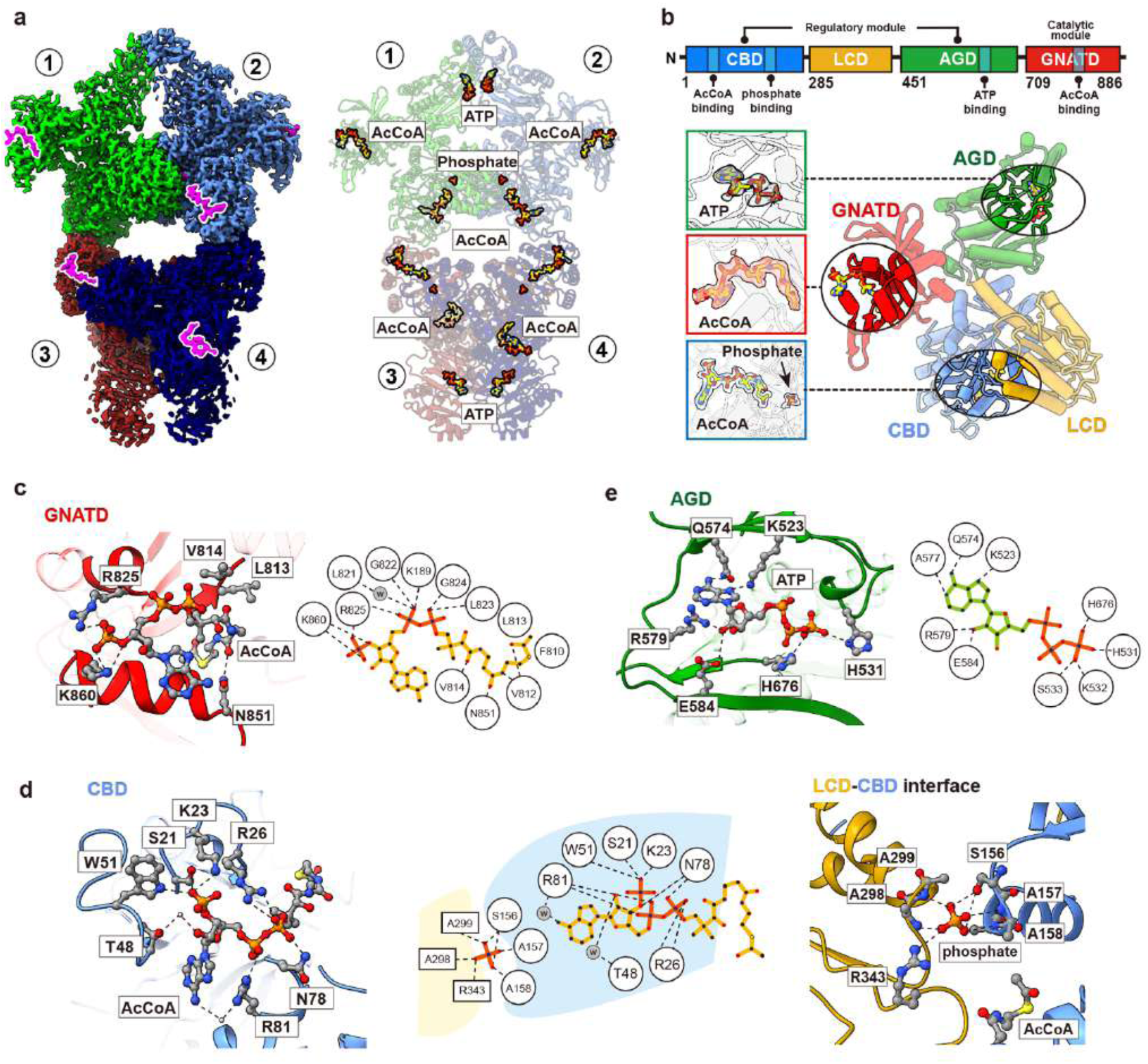
Structure of the liganded-PatZ. **A,** Ligand bindings to tetrameric PatZ and their binding sites. Four ATPs were docked into the tetramer structure based on the ATP bound AGD crystal structure. **B,** PatZ subdomain schematic and its ligand binding site. 2Fo-Fc electron density map of ATP and cryoEM maps of two AcCoAs and phosphate. **C,** Molecular interactions between PatZ GNATD and AcCoA. **D,** Molecular interactions between the axial dimer interface and phosphate, and between PatZ CBD and AcCoA. **E,** Interaction between PatZ AGD and ATP. *’w’ denotes water molecule. Hydrogen bonds and electrostatic interactions are represented by black dashed lines.

#### AcCoA in catalytic GNATD

Within the PatZ subunits, each GNATD adopts a zigzag configuration, oriented outward to maximize its accessibility and functional flexibility. The structural composition of the GNATD includes five α-helices and eight β-strands, forming a donut-shaped tunnel that facilitates the concurrent orientation of AcCoA and the substrate (Supplementary Video 3). This arrangement is pivotal for the enzymatic activity, directing AcCoA and substrate into a productive alignment. The GNATD features a positively charged surface that binds to AcCoA, and a negatively charged surface that assists in positioning acceptor lysine residues. AcCoA is strategically placed within a V-shaped cleft, bordered by β-strands, ensuring precise orientation. The pyrophosphate group of AcCoA is anchored through interactions with the phosphate-coordinating loop (P-loop^38^, G822-G824) including water-mediated hydrogen bonding and a phosphate group on the ribose, which is secured by K860 (Figure 2c). Moreover, the acetate group of AcCoA, crucial for triggering enzymatic reactions, is situated near E809. This residue is identified as a potential catalytic site^32^ and is in close proximity to the putative acceptor lysine binding site, underscoring its significance in the enzymatic process.

#### AcCoA in N-terminal CBD

The PatZ enzyme distinguishes itself through the inclusion of a regulatory domain alongside its catalytic domain, a feature not fully elucidated in terms of its mechanistic role. We visualized a well-defined AcCoA density within the N-terminal CBD (1-285) of PatZ. Notably, the CoA portion of AcCoA is positioned to project outward, while the acetyl group is nestled within the inter-subunit cleft. The binding pocket is characterized by its positive charge and an elongated, narrow morphology, tailored to interact with AcCoA’s three phosphate groups. This interaction is further reinforced by the pocket’s design to accommodate the distinct components of AcCoA through a network of hydrogen bonds including water-mediated interactions (Figure 2d). Particularly, the positively charged residues (K23, R26 and R81) within the pocket engage in electrostatic interactions with AcCoA’s phosphate groups, underscoring the specificity and efficiency of the binding process.

#### Phosphate located in intersubunit interface

We have identified a significant density in the interface between each CBD and LCD derived from different protomers. Considering the homology with ACD1^36^, this density can be attributed to phosphate. The phosphate binding site is located at the tip of the α-helices originating from the LCD and CBD, and is further stabilized by additional loops. The phosphate is spatially adjacent to AcCoA, which resides within the CBD (Figure 2d). Since the phosphate ion is deeply embedded inside the structure, stabilizing a lateral dimer. Of note, this phosphate can be converted to AcP during ATP synthesis pathway for ACD enzyme^39^. Through structural analysis, we found that although PatZ’s regulatory domain is incapable of enzyme catalysis, it still binds phosphate in the same location as ACD1.

#### ATP in ATP grasp domain

The ATP grasp domain (AGD, 451-709) encompasses the ATP binding pocket. Since the dynamic nature of the AGD complicates the visualization of ATP density in cryoEM. Consequently, X-ray crystallography was employed, successfully confirming the presence of ATP within the AGD’s binding pocket at a 2.24 Å resolution. This structure shows that the ATP binding site is nestled in a cleft located in the central AGD, engaging in various interactions with surrounding amino acids (Figure 2e). Despite the presence of 1 mM magnesium during the crystallization process, no electron density map was observed. Instead of a divalent ion, the triphosphate moiety of ATP is stabilized by two histidines (H531, H676) in AGD.

### AcCoA binding activates GNATD

To understand how AcCoA binding activates catalytic GNATD and the chemical basis of its catalytic mechanism, we compared GNATD structures between the apo- and AcCoA-bound states. Binding of AcCoA to GNATD induces significant spatial changes, particularly in the substrate gating motif (V746-T763, hereafter, gating motif), which are located near the substrate access channel and undergo a notable secondary structural transition (Figure 3a, Supplementary video 3). In apo-PatZ, the β-sheet of the gating motif is extended, covering the substrate binding sites of GNATD. However, in the presence of AcCoA, this region transforms into a loop and an α-helix, establishing direct interaction with AcCoA (Figure 3b) and opening a substrate binding pocket (Figure 3c). This structural observation indicates the extended β-sheet of gating motif sterically obstructs the interaction between donor AcCoA and the acceptor substrate. AcCoA binding triggers the rearrangement of the gating motif, removing the physical barrier between AcCoA and the substrate. These structural rearrangements can be crucial to organize the AcCoA donor and the substrate in catalytic proximity. Therefore, we conclude that AcCoA binding is required for GNATD by switching the gating motif into an active structural conformation.

**Figure 3.**
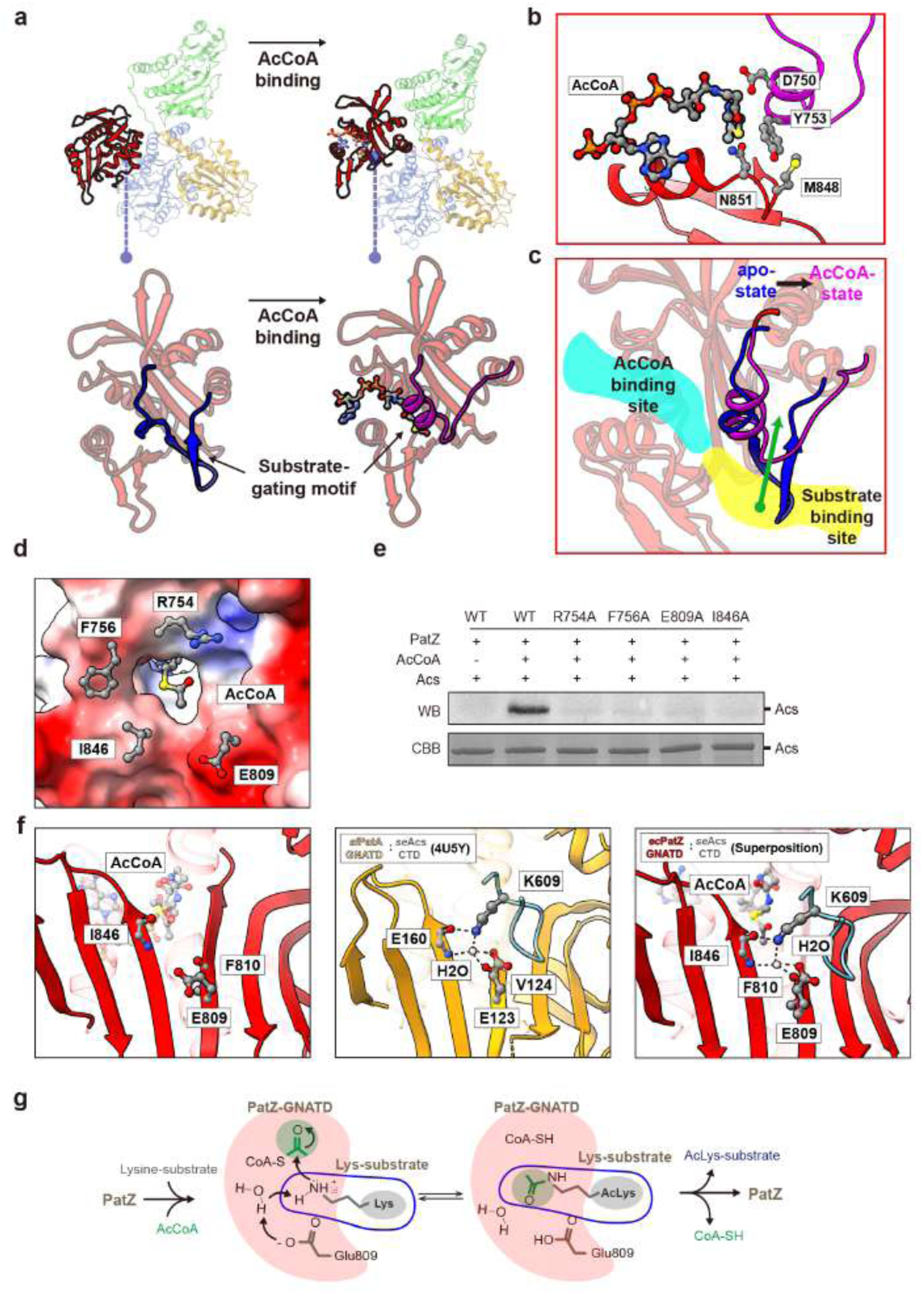
AcCoA-induced conformational change and catalytic residues in GNATD. **A,** Structural rearrangement of GNAT (red cartoon). Substrate gating motifs in the apo-state and AcCoA-bound state are illustrated in blue color and purple color, respectively. AcCoA is shown in ball-and-stick style. **B,** Molecular interaction between AcCoA and PatZ GNATD. Magenta color is part of the substrate gating motif of GNATD. **C,** Structural influence of the AcCoA-induced movement of gating motif on the formation of the substrate binding site. **D,** Electrostatic surface charge and amino acid residues comprising the acceptor lysine binding site in GNATD. **E,** Comparison of relative functional level between wild-type and point mutants which comprise acceptor lysine binding pocket. Western blot analysis was performed using an anti-acetylated lysine antibody. **F,** Homology structure based catalytic mechanism prediction of PatZ using *sl*PatA and seAcs complex structure. Hydrogen bonds are represented by black dashed lines. **G,** Proposed acetyltransferase mechanism of PatZ.

Based on the active conformation of GNATD, we investigated the critical residues that potentially contribute to enzyme activity and substrate binding. By focusing on the distinctive donut hole shape of the GNATD, we observed that the acetyl group of the acetyl-donor AcCoA is positioned at the center of this hole (Figure 3d), oriented towards the substrate binding site. This region contains several charged residues, and we created alanine mutants for four key amino acids (R754, F756, E809, I846) and compared their enzyme activity to that of the wild-type. The results showed that mutations in any of these four residues led to a significant reduction in enzyme activity (Figure 3e). Based on these functional tests and structural analysis, we confirmed that the acetyl group of AcCoA and the acetyl-acceptor lysine are strategically positioned around the donut hole of the GNATD, playing a crucial role in the acetyl-transfer reaction.

To gain a deeper chemical understanding of the enzyme mechanism, we compared our structure with that of the GNATD of *Streptomyces lividans* PatA (*sl*PatA), which has been structurally characterized in complex with *S. enterica* Acs (*se*Acs) (PDB ID: 4U5Y)^40^. The GNATD of *sl*PatA is structurally similar to that of PatZ GNATD, with an RMSD of 1.12 Å (Extended Data Figure 3a). By integrating structural information, we identified the arrangement of key catalytic residues in PatZ, including E809 and the carbonyl oxygens of F810 and I846 (Figure 3f). Notably, these residues are highly conserved within the GNAT family. E809, in particular, is recognized as a crucial catalytic residue (Extended Data Figure 3b). Of note, this glutamate residue can activate a water molecule, which coordinated by E123^PatA^, carbonyl oxygen of V124^PatA^, amide nitrogen of E160 and K609^Acs^, to remove a proton from the lysine amine group, facilitating a nucleophilic attack on the carbonyl carbon of enzyme-bound AcCoA (Figure 3f). Taken together, the structural similarities between *sl*PatA and *E. coli* PatZ, coupled with the conserved nature of the catalytic residues, underscore the mechanistic insights into the enzymatic function of PatZ (Figure 3g). These findings imply the critical roles of E809 and the amino acid backbones in facilitating the acetyl-transfer reaction, advancing our understanding of the GNAT family’s catalytic mechanisms.

### Allosteric regulation for PatZ

Next, to investigate the functional and mechanistic association between the regulatory domain and catalytic GNATD, we generated domain-wise truncated PatZ mutants and examined their activity. The AGD-truncated mutant (trAGD) produced soluble aggregates, while the other truncated mutants (trCBD-LCD, GNATD-only and regulatory domain-only (RD-only)) were properly folded as evidenced by size exclusion chromatography (Figure 4a). As expected from our structural data, the RD-only mutant formed a tetramer, whereas the trCBD-LCD and GNATD-only mutants existed as monomers based on their elution profiles. As anticipated, the RD-only mutant without catalytic module abolished enzymatic activity. Interestingly, both the GNATD-only mutant and the domain-wise truncation mutants of the regulatory domains also lost their acetyltransferase activity towards Acs (Figure 4b). It is noteworthy that the trAGD mutant, which showed aggregation, likely experienced a loss of activity related to the instability of its truncated structure (Extended Data Figure 4a). The loss of enzymatic activity in both the GNATD-only and truncated regulatory domain mutants suggests the necessity of oligomeric conformation and highlights the critical interplay between these domains for the acetyltransferase function of PatZ.

**Figure 4.**
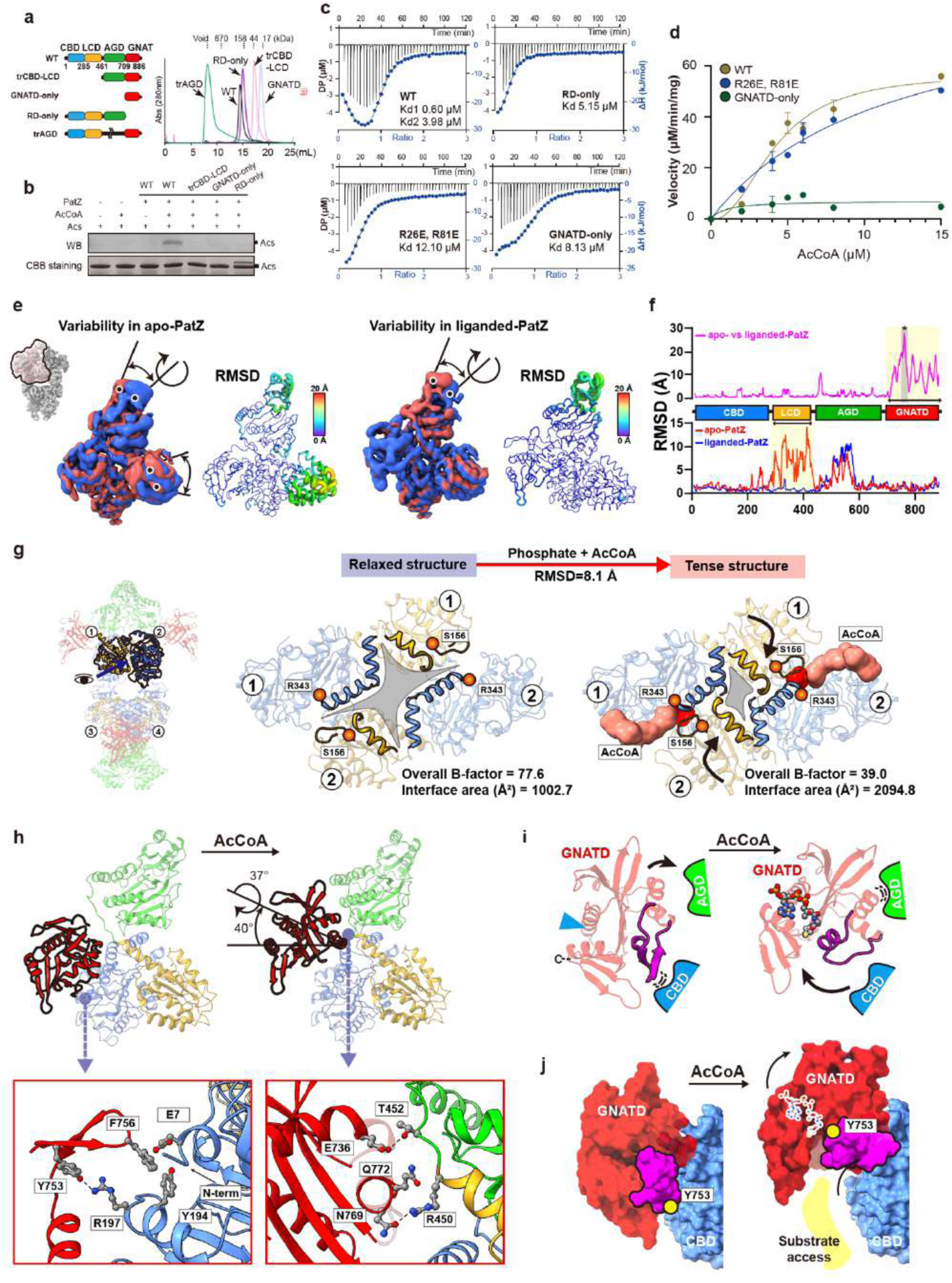
Allosteric coupling of regulatory domain to GNATD. **A,** Schematic of domain truncation construct design and its size exclusion chromatography peak pattern. ‘tr’ denotes truncation. **B,** Comparative enzymatic activity analysis of subdomain truncated constructs via western blot. Western blot analysis was performed using an anti-acetylated lysine antibody **c,** ITC profiles of AcCoA binding to wild-type, truncated forms and RR (R26E/R81E) mutant of PatZ. **D,** Comparative enzymatic activity analysis of wild-type, RR mutant and GNAT-only constructs of PatZ. **E,** Structural variability comparison between apo-(left) and liganded-PatZ (right). **F,** Comparison of backbone RMSD between apo- and liganded-PatZ (purple) and backbone RMSD within apo-PatZ variability (red) and liganded-PatZ variability (blue). Asterisk refers to the substrate gating motif of GNATD. **G,** Ligand binding-mediated structural change of PatZ regulatory domain **h,** Structural effect of AcCoA binding on the interaction between GNATD and regulatory domain. Hydrogen bonds are represented by dashed lines. **I,** Role of substrate gating motif in changes in interaction between GNATD and regulatory domain following AcCoA binding **j,** Substrate access channel formed by AcCoA-mediated GNATD conformational change and the role of substrate gating motif. Magenta color is a space filling model of substrate gating motif.

To investigate whether and how AcCoA binding to the regulatory domain affects PatZ activity, we analyzed the AcCoA binding affinity of PatZ using isothermal titration calorimetry (ITC). As anticipated from a previous study on *S. enterica* Pat^31^ and our structural data in this study (Figures 2 and 3), AcCoA binding to wild-type PatZ exhibited a biphasic curve, best fitting to a two-site binding model. One binding site demonstrated a much stronger affinity (K_d1_ = 0.60 μM) compared to the other (K_d2_ = 3.98 μM) (Figure 4c). However, when residues R26 and R81 within the AcCoA-binding pocket of PatZ (see Figure 2d) was mutated to Glu, disabling AcCoA binding to the RD, the resulting RR mutant of PatZ showed one-site binding (K_d_ = 12.10 μM), similar to, but with lower affinity than, the RD-only form (K_d_ = 5.15 μM). These kinetic data suggest that the RD has a higher AcCoA binding affinity than the GNATD and that binding of AcCoA to one site increases the affinity of the other site, indicating positive cooperativity. Notably, the monomeric GNATD-only form exhibited a binding affinity (K_d_ = 8.13 μM) similar to the tetrameric RR mutant, indicating that the AcCoA binding affinity of GNATD is not significantly affected by oligomerization.

Next, we conducted the PatZ activity assay by measuring the free sulfhydryl group of CoA released after the acetyltransferase reaction. The formation of 2-nitro-5-thiobenzoate (TNB2-) from the reaction between the free sulfhydryl group and DTNB (5,5’-dithio-bis-(2-nitrobenzoic acid)) was recorded at 412 nm. Consistent with previous findings for *ec*PatZ^32^ (Hill coefficient = 7.91) and *se*PatZ^31^ (Hill coefficient = 2.2), wild-type PatZ exhibited a sigmoidal activity curve, indicative of positive cooperativity (Hill coefficient = 2.21; K_half_ = 4.20 μM) (Figure 4d). However, the RR mutant exhibited a typical Michaelis-Menten curve and had a significantly lower AcCoA affinity (K_m_ = 11.78 μM) compared to wild-type PatZ. These results confirm that AcCoA binding to the regulatory domain not only increases GNATD’s affinity for AcCoA but is also essential for the proper allosteric regulation of PatZ’s enzymatic activity.

To understand the mechanistic relationship with AcCoA-binding between the regulatory domains and GNATD, we analyzed the domain-wise structural dynamics of PatZ upon ligand binding using single protomer heterogeneity from our cryoEM images. PatZ displayed varying degrees of structural dynamics with and without AcCoA (Figure 4e, Supplementary Video 4). Specifically, apo-PatZ exhibited significantly higher structural heterogeneity compared to the ligand-bound form, as also evidenced by resolution improvement from 2.56 Å to 1.99 Å. The AGD’s motion was less correlated and continuous regardless of AcCoA binding. However, we identified three distinct classes (class I-III, Extended Data Figure 4b) of domain interfaces of the LCD with ∼15 Å RMSD in apo-PatZ, while the AcCoA-bound structure showed a well-converged single conformation (Figure 4f). These structural changes in apo-PatZ resulted in a relaxed axial intersubunit space between the LCD of one subunit and the CBD of the neighboring subunit. To characterize changes in the interface, we measured the distance between S156 and R343 of the neighboring protomer, forming part of the ligand binding pocket (Figure 4g, Extended Data Figure 4c). In the apo-state, this distance expanded to 16.9 Å. In the ligand bound form, the distance converged to 11.9 Å. Notably, the class III map in apo-PatZ displayed attributable density to phosphate with an RMSD of 1.2 Å compared to the AcCoA-bound conformation. Therefore, our analysis indicates that AcCoA binding in the CBD contributes to stabilizing the complex and enforcing a stably closed intersubunit conformation.

Interestingly, the GNATD itself displays well-defined static structures in both the apo- and ligand-bound states. However, binding of AcCoA to PatZ induces significant domain-wise movement in GNATD, specifically a ∼37° rotation and a ∼40° tilt relative to the regulatory domain from the apo- to AcCoA-bound conformation (Figure 4h, Supplementary Video 4). We identified that the regulatory domains alter their specific interactions with the GNATD upon ligand-induced conformational transition. In apo-PatZ, the N-terminal end of CBD (Figure 4i) interacts with the β-sheet of gating motif, anchoring GNATD in a down-position. Conversely, in the ligand-bound conformation, the AGD and the interdomain loop (E451-S561) establish new specific interactions with the hinge region (E736, Q772, N769) of GNATD, stabilizing GNATD in an up-position (Figure 4h).

Importantly, in the apo-PatZ, the tethered configuration limits the space between GNATD and CBD, resulting in the closure of the substrate access channel, with the β-sheet of gating motif physically obstructing the interaction between donor AcCoA and the substrate (Figure 4j, left panel). Binding of AcCoA to both the CBD and GNATD opens the substrate access channel and triggers the rearrangement of the gating motif into an active conformation (Figure 4j, right panel). Taken together, the regulatory domain is essential for the activation of GNATD, and AcCoA binding to both the CBD and GNATD positively cooperates in allosteric regulation to facilitate PatZ’s enzymatic reaction, which is in accordance with the ITC and DTNB results (Figure 4c,d). Notably, considering that the close conformation of the regulatory domain (class III in apo-PatZ) still displays a down-position of GNATD, it is logical to conclude that AcCoA binding to GNATD and the switching interaction from the regulatory domain are both prerequisite events for PatZ activation.

### Regulatory domain directly shapes Acs binding pocket, enhancing substrate interaction

Upon elucidating a cooperative mechanism between the GNATD and regulatory domains of PatZ for ligand-induced substrate gating, we extended our investigation to explore whether the regulatory domain directly influences binding or specificity of a protein substrate. AlphaFold^41^ predicted the reliable models showing monomeric interaction within the PatZ-Acs complex structure (Extended Data Figure 5). These predictions illustrated that Acs interacts with PatZ and notably, the Acs K609 residue, a known acetylation site, is oriented towards GNATD (Figure 5a) and is positioned within the putative acceptor lysine binding pocket, which is consistent with our experimental analysis (Figure 5a,b). This intricate interaction of Acs is established by an orchestration with the CBD, LCD and GNATD of PatZ.

**Figure 5.**
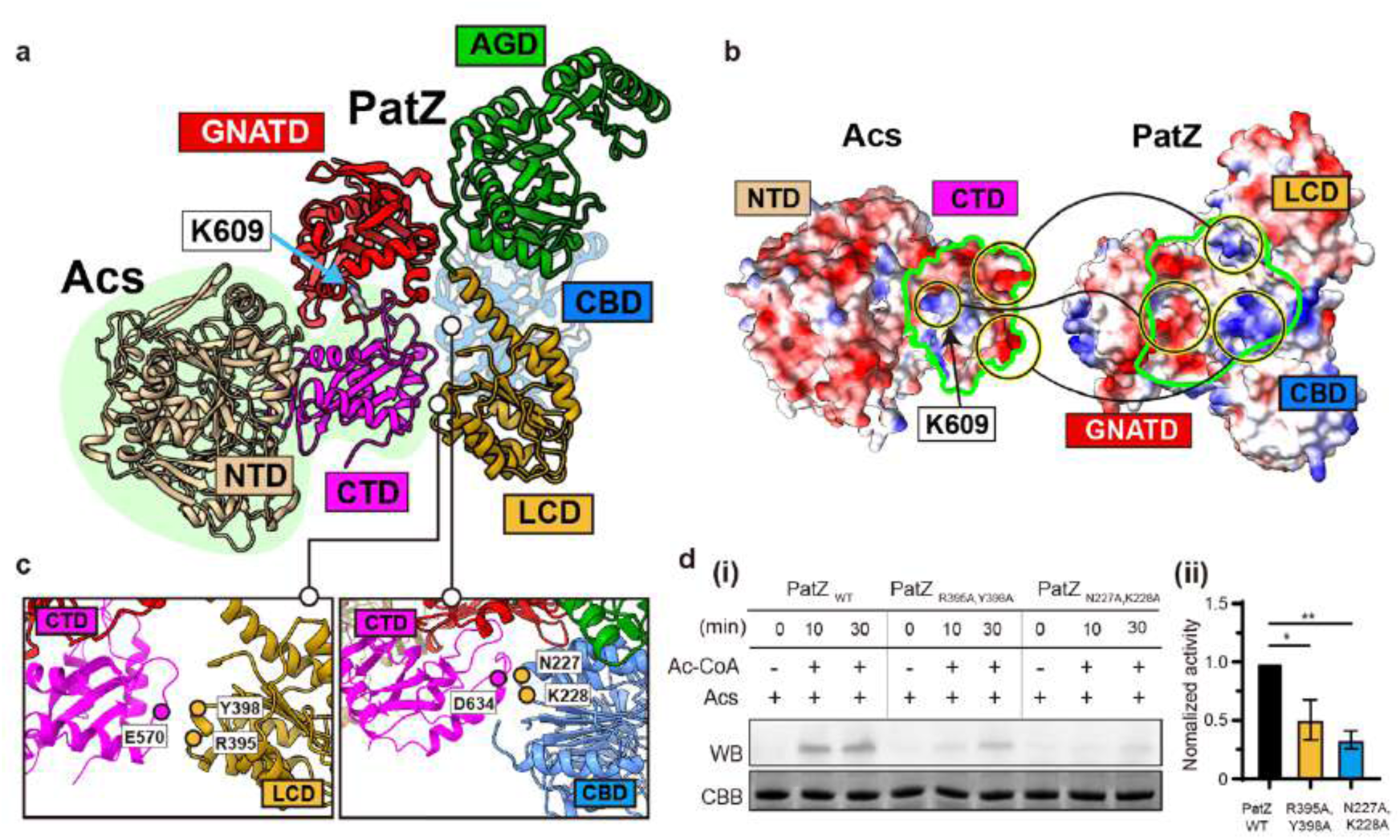
The regulatory domain of PatZ contributes to the formation of Acs binding pocket. **a,** Predicted structure of the *E.coli* PatZ-*E.coli* Acs complex using AlphaFold^41^. Each domain is represented in a different color using cartoon-style representation. Acs K609 is represented in stick style. **b,** Electrostatic surface potentials of Acs and PatZ. Red and blue for negative and positive charges, respectively, and white color represents the neutral area. The circularly linked different parts represent the interacting components. **c,** Acs-CTD and PatZ-LCD interaction (left panel) and Acs-CTD and PatZ-CBD interaction (right panel). The circled regions denote the positions of the labeled residues. **d,** Time-dependent comparison of PatZ mutant activity relative to wild-type. Western blot analysis using anti-acetylated lysine antibody (left panel) and normalized activity comparison graph (right panel).

Significantly, the predicted PatZ-Acs structure underscores that the regulatory domain forms a distinctive groove and binding surface, accommodating the C-terminal domain of Acs and fostering highly complementary polar interactions (Figure 5b). These specific residues exhibit robust coupled variance, indicative of residue-level coevolution between Acs and the regulatory domain of PatZ. This coevolution suggests a finely tuned evolutionary adaptation, underscoring the additional importance of the regulatory domain for PatZ’s functional integrity.

To validate the impact of regulatory domain and Acs interaction on enzyme activity, we identified and mutated coevolved residues at the interface between Acs-CTD and the regulatory domain of PatZ. We engineered R395A/Y398A and N227A/K228A mutants of PatZ (Figure 5c) and evaluated their enzyme activity compared to the wild-type over time. Our results disclosed a pronounced decrease in enzyme activity for both mutants (Figure 5d), signifying the critical role these residues play in maintaining enzymatic function. This significant reduction in activity underscores the importance of the regulatory domain in facilitating proper substrate binding and, consequently, effective catalysis.

## DISCUSSION

We characterized multiple ligand-PatZ interactions and demonstrated that ligand-induced conformational changes facilitate the allosteric regulation of the catalytic activity (Figure 4, 6a). Our investigation delineates PatZ’s working mechanism and highlights key aspects of how ligand binding influences PatZ activity and regulation (Figure 6a).

**Figure 6.**
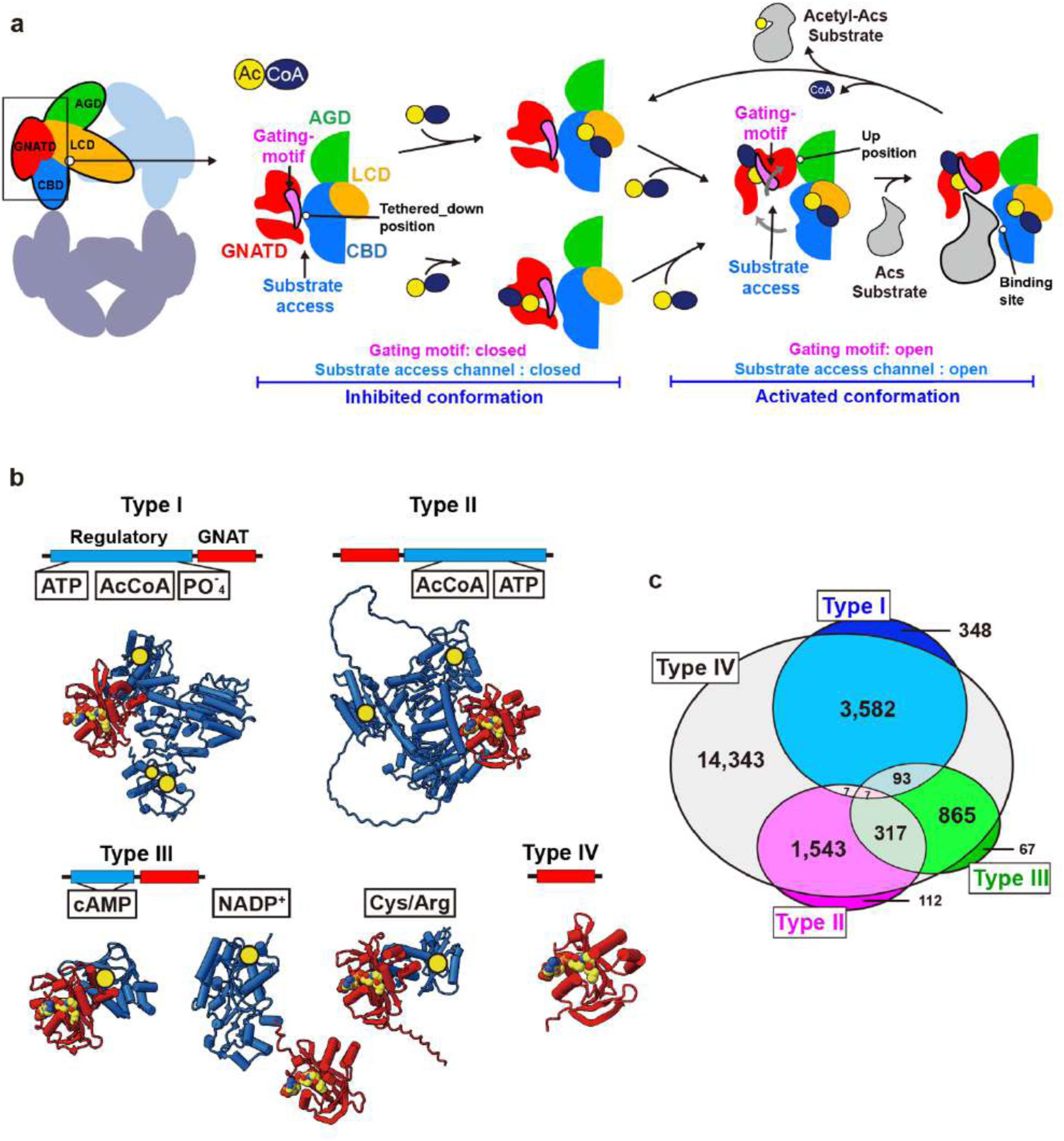
Phylogenetic analysis and catalytic mechanism. **a**, Schematic illustration of ligand-induced domain-specific conformational changes regulating PatZ catalytic activation. **b**, Visualization of Type I, Type II, cAMP-binding type III, ACT type III, NADP+-binding Type III and Type IV **c**, Venn diagram showing the overlap of GNAT types. Each number represents the count of bacterial species containing the corresponding GNAT type.

### Gating motif contributes to auto-regulation of GNATs

Located near the substrate binding site in GNATD, the gating motif (V746-T763) undergoes significant structural changes, demonstrating the mechanism of opening and closing a passage that connects the donor AcCoA and acceptor protein substrate (Figure 4h-j, 5). In the absence of AcCoA, the gating motif sterically occludes substrate binding sites and masks the passage, indicating inhibited conformation (Figure 4h-j). Upon AcCoA binding, the gating motif coordinates with the AcCoA to expose the substrate binding sites and aligns the proximity between AcCoA and protein substrates, indicating activated conformation (Figure 4h-j). AcCoA induced-structural changes in GNATs have been previously discussed in the studies of RimL from *S. typhimurium*^42^, Rv0819 from *M. tuberculosis*^43^ and *ss*Pat from *Sulfolobus solfataricus*^44^. RimL and *ss*Pat are GNATD only enzymes without regulatory domain (Type IV GNAT), while Rv0819 is a tandem-repeated GNAT enzyme (Type V GNATs). In these studies, the homologous region with a gating motif is referred to as α1-α2 loop, mobile loop or bent helix, which remain unresolved in the absence of AcCoA due to their intrinsic dynamic features. This loop becomes rigidly ordered by AcCoA binding, resulting in the formation of a substrate binding pocket, similar to the gating motif of PatZ. Notably, the mutation study of *ss*Pat has shown that the bent helix is essential for enzyme activity.

Importantly, this structural component, the gating motif in GNATs, is not only present in prokaryotes but also preserved in eukaryotes, including humans (Extended Data Figure 7), which suggests a common mechanism of GNATs’ auto-regulation across species. Collectively, based on the clear visualization of the inhibitory and active conformations of GNATD in PatZ, we propose an auto-regulatory mechanism of GNATs mediated by the gating motif. Interestingly, despite the high structural similarity of GNATs, the gating motif exhibits relatively low sequence similarity^45^ (Extended Data Figure 7). Since the gating motif also contributes to the formation of the substrate binding pocket, it is tempting to speculate that the gating motif may be involved in substrate specificity, which requires further studies.

### Regulatory domain interaction mechanistically regulates PatZ activity

We characterized that ligand binding to the regulatory domain significantly stabilizes intersubunit interface and rigidifies PatZ’s homotetrameric architecture (Figure 4g). Concurrently, this conformational stabilization of the regulatory domain is correlated with the activation process in the gating motif induced by AcCoA binding to GNATD (Figure 4i). These cooperative structural events result in the domain-wise conformations of the up- and down-positioning of the GNATD. The gating motif of GNATD interacts with the CBD in the absence of AcCoA, leading to the down-positioning of GNATD (Figure 4i, left panel). In this down position, the gating motif tethered by the regulatory domain illustrates a restricted space for the substrate access channel (Figure 4j, left panel), indicating an inhibitory conformation. Upon AcCoA binding, the gating motif loses its interaction with CBD, allowing GNATD to refresh its interaction with AGD and adopt an up-positioned conformation (Figure 4i, right panel). This position creates a widely exposed space between GNATD and CBD, allowing protein substrates access to the substrate binding site (Figure 4j, right panel), indicating an activated conformation.

Interestingly, the stabilized structure with phosphate bound PatZ (Extended Data Figure 4b,c) displays the down position of GNATD and the inactive conformation of the gating motif. The regulatory domain of this structure is identical to that of the AcCoA-bound structure, implying that the stability of the regulatory domain alone cannot transform GNATD into its active conformation. GNATD alone does not also have enzymatic activity and thus suggest that AcCoA binding to both CBD and GNATD is a prerequisite for activating PatZ. Importantly, we (Figure 4c,d) and others^31^ have demonstrated the cooperative relationship of AcCoA binding between GNATD and CBD. AcCoA binding to the wild-type GNATD is approximately 2 times higher than to the GNATD-only protein and 6.6 times higher to the regulatory domain than to GNATD (Figure 4c). These findings collectively suggest that AcCoA binding to CBD reduces the dynamics of the regulatory domain, enhances the AcCoA binding to GNATD, facilitates the rearrangement of the gating motif, and activates the enzyme.

Therefore, our study elucidates the active and inactive conformations in the cooperation between GNATD and the regulatory domain and highlights PatZ’s activity regulation mechanism. This elegant mechanism employs a dual strategy that switches between active and inactive conformations and allosterically modulates enzyme activity upon cooperative AcCoA binding.

### Regulatory domain contributes PatZ’s substrate specificity

Prior studies have highlighted the unique specificity of PatZ for Acs^32,46^, and it has been also observed that strains lacking PatZ do not produce acetylated Acs^47^. Through our observations, we identified a structural opening event in GNATD, in cooperation with CBD, that forms an Acs-specific binding pocket (Figure 5). These findings validate additional functional layers of the regulatory domain. Moreover, it is evident that GNATD alone in PatZ lacks enzymatic activity toward Acs^40^ (Figure 4b), and point mutations in the regulatory domain that disrupt Acs interaction significantly reduce Acs acetylation (Figure 5c,d). Our results further demonstrate that Acs binds complementary to the substrate binding site formed by both GNATD and the regulatory domain (Figure 5a,b). This is supported by the observation that *E. coli*-derived YiaC, which lacks a regulatory domain (GNAT type IV), is inactive to Acs (Supplementary Data Figure 8). The residues involved in Acs-PatZ interaction are highly conserved, suggesting a finely tuned evolutionary adaptation that underscores the regulatory domain’s critical importance for PatZ’s function concerning Acs.

While we elucidate substrate-specific mechanisms, it is noteworthy that PatZ is recognized for its ability to acetylate a diverse array of protein substrates within the bacterial proteome, with an estimated around 1000 substrates compared to around 20∼600 for GNATD-only enzymes like YjaB, YiaC, RimI and PhnO^48^. Although these acetyl-substrate repertoires overlap significantly, PatZ exhibits a unique capability for the specific targeting of substrates^32,46^.

Our study implies that PatZ employs dual modes of substrate recognition: a general acetyltransferase function for exposed lysines across a broad spectrum of proteins and a specific mechanism through cooperative substrate recognition involving both GNATD and the regulatory domains (Figure 6a). This dual functionality highlights the sophisticated regulatory mechanisms that enable PatZ to achieve both substrate specificity and enzymatic versatility.

### Diverse roles of regulatory domains in GNATs family for bacterial physiology

Protein acetylation plays crucial roles in cellular physiology by impacting various processes and regulatory mechanisms^49^. Present in all living organisms across the Bacteria, Archaea, and Eukarya domains^50^, protein acetyltransferases, including GNATs suggest that protein acetylation has existed since the earliest stages of life, predating the Last Universal Common Ancestor (LUCA) more than 4 billion years ago^51^. Despite the vast evolutionary divergence from LUCA, essential cellular functions such as genetic information processing and core metabolic pathways have been conserved across phylogeny^52^.

Our study reveals that the mechanistic coupling of AcCoA binding to the regulatory domain modulates the activity of the Type I GNATs, such as PatZ. Given that unique types of regulatory domains in other GNAT’s can bind different ligands, this variation suggests that bacteria regulate GNATs’ activity in response to diverse metabolic conditions (Figure 6b). We conducted a protein structure-based phylogenetic analysis against the AlphaFold Protein Structure Database^53^ to explore the distribution of GNAT types in bacteria (Extended Data Figure 6). Notably, we found that while bacteria can simultaneously possess multiple types of GNATs, Type IV GNATs, which consist solely of the GNATD, exhibit a ubiquitous distribution across the bacterial domain (Extended Data Figure 6e,f). In contrast, we identified a distinct pattern where other types of GNATs with unique regulatory domains are predominantly found within specific phyla (Extended Data Figures 6e,f).

Type IV GNATs, processing a single catalytic GNAT domain, are ubiquitously present in bacteria (Extended Data Figure 6e,f), where they regulate nucleoid organization^54^, transcription^55^, and central metabolic pathways^56^. This ubiquity implies GNATs’ involvement in regulating housekeeping processes. In contrast, Type I, II, and III GNATs possess additional regulatory domains (Figure 6b, Extended Data Figure 6b-d), likely enabling them to sense various environmental conditions and tightly regulate genetic information flow and metabolic fluxes in response to nutrient availability and cellular energy status.

For instance, in *E. coli*, the Type I GNAT PatZ uses AcCoA not only as an acetyl-donating substrate but also as a ligand binding to its regulatory domain (Figure 2), thus allosterically regulating its GNAT activity (Figure 4). One of the most studied Type II GNATs is PatA from *S. lividans*^57^. While the ligands binding to its regulatory domain are not well understood, predictions using AlphaFold^41^ and AlphaFill^58^ revealed that AcCoA and ATP bind to PatA’s regulatory domain (Figure 6b). Previous studies have reported that Type III GNATs have their activity regulated by the binding of amino acids, cAMP, and NADP^+^ to their regulatory domains^59–61^. Although different types of GNATs adopted various ligands as regulatory cues, these ligands generally represent nutritional and cellular energy status^62^.

Interestingly, the activity of Acs is regulated by acetylation via GNATs in most, if not all, bacterial species studied so far^4^. AcCoA is a crucial metabolite in central metabolism, acting as a crossroad in pathways such as glycolysis, gluconeogenesis, the tricarboxylic acid cycle, the glyoxylate bypass, lipid metabolism, and amino acid synthesis^63^. Nearly all life forms have evolved to adopt AcCoA as a key metabolic intermediate, making it essential to maintain its levels for supporting all cellular processes. While different bacterial phyla have adopted different types of structures of regulatory domains (Figure 6b-d), a common function of GNATs appears to be the regulation of housekeeping processes by sensing diverse cellular cues.

## MATERIAL AND METHODS

### Constructs design and molecular cloning

The full-length or truncated PatZ gene (UniProt ID: P76594) was cloned into the pETDuet-1 vector (Novagen) with a 6xHis tag at the C-terminus. The full-length Acs gene (UniProt ID: P27550) was cloned into the pETDuet-1 vector with an N-terminal 6xHis tag. All mutants were generated by site-directed mutagenesis.

### Protein expression and purification

The recombinant DNA is transformed into the *patz, pta* deletion ER2566 strain. The transformed cells were grown in LB medium at 37 °C until the O.D 600 value reached 0.6∼0.8. The temperature was then lowered to 17 °C, and 1 mM IPTG was added to induce protein expression. After 16 hours of induction, the cells were harvested by centrifugation and the medium was completely discarded for storage at - 80 °C. The harvested cells were disrupted using Emulsiflex C3 (Avestin) and the cell debris was removed by centrifugation at 20,784 g for 30 min. The supernatant is filtered before purification by FPLC. And the filtered samples were purified by Ni-NTA affinity chromatography using Histrap HP pre-packed columns (Cytiva). After washing with wash buffer (25 mM Tris-HCl pH 7.4, 500 mM NaCl, 20 mM imidazole, 1 mM MgCl_2_, 10% glycerol) elution was performed with elution buffer (20 mM HEPES pH 7.4, 500 mM NaCl, 1 mM MgCl_2_, 10% glycerol, 1 mM DTT) with 300 mM Imidazole. The purified protein samples were then further purified by size exclusion chromatography (SEC) using a Superose 6 Increase column (Cytiva) in a buffer containing PBS with additional 100 mM NaCl. Acs was cultured in TB medium under the same conditions. The purification procedure followed the same protocol: After extensive washing with wash buffer (25 mM Tris-HCl pH 7.4, 500 mM NaCl, 40 mM imidazole, 10% glycerol), elution was carried out using elution buffer (25 mM Tris-HCl pH 7.4, 500 mM NaCl, 300 mM imidazole, 10% glycerol). and then further purified by SEC using a HiLoad 16/600 superose 6 pg (Cytiva) in a buffer containing 25 mM Tris-HCl pH 7.4, 500 mM NaCl, 5% glycerol.

### CryoEM specimen preparation and imaging condition

Purified protein sample was adjusted to 1.5 mg/mL and vitrification was conducted using the Vitrobot system (ThermoFisher). 3 μL samples were applied to the discharged UltrAuFoil R1.2/1.3 gold 300 mesh (Quantifoil). Grids were blotted for 3 sec and plunged into liquid ethane at 100% humidity at 8 °C. The grids were roughly screened on a Glacios operated at 200 keV (cryoTEM, ThermoFisher) and equipped with a Falcon 4 direct electron detector. The nicely vitrified grids were placed on a 300 keV Titan Krios G4 (ThermoFisher) equipped with a Falcon 4i direct electron detector. Data was acquired in EER format at a pixel size of 0.723 Å/pixel using aberration-free image shift (AFIS) to accelerate the data acquisition. and total exposure dose was 50 e/A^2^ with -0.8 to -1.5 μm defocus range. More details about imaging conditions are described in Supplementary Table 1.

### CryoEM image processing

The EM data processing workflow for all datasets is illustrated in **Extended Data Figures 1 and 2**. The processing was conducted using CryoSPARC v4.4.1 packages, and a comparable strategy was employed for all datasets. Firstly, the beam-induced motion of the movies was corrected using Patch motion correction, and the contrast transfer function (CTF) of each micrograph was calculated using Patch CTF estimation. Some micrographs were excluded due to insufficient resolution of the CTF and defocus range. Subsequently, about 3,000 particles were selected via blob-based picking from a subset of 30 micrographs, resulting in the generation of 2D templates. Finally, a considerable number of particles were automatically selected from the entirety of the micrographs using the templates. The extraction of particles was conducted using a box size of 448 X 448 pixels. Subsequently, several iterations of 2D classification were performed, and 3D volumes were generated without alignment through Ab-initio reconstruction. The optimal volume class with D2 symmetry was subjected to Homogeneous 3D refinement. To enhance the overall quality of the particles and volumes, Topaz particle picking and Reference-based motion correction (beta) were applied based on the initially refined map. Subsequently, Final maps were generated, followed by global and local CTF correction to enhance the map.

For asymmetric unit analysis, each particle was expanded according to D2 symmetry. The optimal volumes for each domain and subunit were classified through multiple rounds of 3D classification and subjected to local refinement using the corresponding masks without any symmetry enforcement. These refinements significantly improved domain resolution and features. Structural heterogeneity was characterized by iterative 3D classification and 3D variability analysis. To generate representative final maps, locally refined maps were aligned into the consensus refined map and patched using the “vop maximum” command in UCSF ChimeraX^64^. These focused refinement maps and the merged maps were subsequently used for model building, refinement, and analysis.

### Model building and refinement

Initial models were generated using AlphaFold and subsequently fitted to the map using ISOLDE. Manual model fitting was then performed using the Coot program, followed by model refinement using Real-space refinement in Phenix. The refinement results and model quality were numerically assessed through Real-space refinement output values and MolProbity scores. The structure of liganded-PatZ was built using the structure obtained from apo-PatZ as a template, and model refinement was carried out following the same methodology as for apo-PatZ.

### X-ray crystallography, data processing and refinement

AGD of PatZ (T463-S695) was subcloned into pET28a and overexpression and purification were conducted using the same method but with an SEC buffer containing 20 mM HEPES-NaOH pH 7.5, 150 mM NaCl, 1 mM MgCl_2_, 1 mM DTT. Concentration of purified samples was adjusted to 10 mg/mL and co-crystallized with 5 mM ATP. Initial crystallization screening was conducted by using a Mosquito robot (TTP Labtech) and a single, appropriate size of crystals appeared at 0.2 M potassium sodium tartrate, 20 % (w/v) PEG 3350. 20 % ethylene glycol was added as a cryo-protectant and flash frozen in liquid nitrogen. Crystal diffraction was carried out at the Pohang accelerator laboratory 5C (PAL-5C) beamline (wavelength= 0.97942 Å)^65^ on 100K temperature. Diffraction data sets were processed and scaled with the programs HKL2000^66^ and imosfilm^67^. The phasing information was solved by a molecular replacement (MR) method using the cryoEM structure. MOLREP^68^, REFMAC5^69^, and COOT were used for MR, structure refinement and further modeling, respectively. Figures were prepared using PYMOL and ChimeraX. The Ramachandran plot analysis of the final model revealed that 96.44% of residues were in favored regions, 3.34% in allowed regions, and 0.22% in disallowed regions.

### In vitro acetylation assay: Western blot

Acs (1.3 μM) was incubated with wild-type or mutant PatZ (425 nM) in the presence of 1 mM AcCoA (Cayman) or 1-10 mM of AcP (Merck) in an assay buffer (25 mM sodium phosphate pH 8.0, 0.5 mM EDTA, 100 mM NaCl) at 37 °C for 10 min, if not otherwise specified. Proteins were then separated by 4-20% gradient sodium dodecyl sulfate polyacrylamide gel electrophoresis (SDS-PAGE) and transferred onto a nitrocellulose membrane in transfer buffer for 70 min at 200 mA at 4°C. After transfer, membranes were blocked with 4.5% BSA in PBS-T (4 g NaCl, 100 mg KCl, 720 mg Na_2_HPO_4_, 123 mg KH_2_PO_4_, 250 μl Tween-20 in 500 ml) for 60 min at room temperature. After blocking, the membrane was washed using PBS-T once and then incubated overnight at 4°C with the first antibody, Anti-acetyl lysine antibody (ICP0380, Immunechem), diluted 1:5000 in PBS-T containing 4.5% BSA. After the treatment with the first antibody, the membrane was washed three times with PBS-T for 10 minutes each. Then, it was incubated with goat anti-rabbit IgG-horseradish peroxidase (HRP)linked secondary antibody (LF-SA8002, Abfrontier), diluted 1:5000 in PBS-T containing 4.5% BSA, for 60 min. The membrane was washed 4 times with PBS-T for 10 min each time, incubated in Western HRP substrate (WBLUF0100, Millipore), and imaged in the Luminograph II (ATTO)^70^.

### In vitro acetylation assay: DTNB assay

The reaction mixture was prepared by adding wild-type or mutant PatZ (1 μM), Acs protein (20 μM), DTNB (100 μM, 5,5’-dithio-bis-(2-Nitrobenzoic Acid, Thermo), and AcCoA (from 0 to 20 μM, Cayman) in a 200 μL volume of reaction buffer (25 mM sodium phosphate, pH 8.0, 0.5 mM EDTA, 100 mM NaCl). Given that both Acs and AcCoA serve as substrates for PatZ, experiments utilized lower concentrations of AcCoA compared to Acs to obtain an accurate initial velocity of CoA release. Absorbance at 412 nm wavelength was monitored every 5 seconds for 10 minutes at 37°C using a Flex3 microplate reader (Molecular Devices). A standard curve was created using CoA (Avanti) concentrations ranging from 0 to 100 μM. Velocity was determined by measuring the consumption of AcCoA using the absorbance at 412 nm and then converting this value using the CoA standard curve. This consumption rate was then divided by the time (in minutes) and the amount of PatZs in mg. Velocity and AcCoA concentration were used to fit the wild-type PatZ to a sigmoidal allosteric curve, while the mutants were fitted to a Michaelis-Menten curve, using Prism (GraphPad) analytical software, as used in a previous study^31^.

### Isothermal Titration Calorimetry (ITC) assay

The protein sample was prepared in the ITC buffer (1X PBS, 100 mM NaCl), and AcCoA was prepared in the same ITC buffer at a 15-fold concentration of each protein. For titration, samples were loaded into a 96-deep-well plate three consecutive wells filled for a single titration: samples for the reaction cell, the injection syringe, and the binding buffer for pre-rinsing the reaction cell. All titrations were performed using the automated MicroCal AutoPEAQ ITC instrument (Malvern Panalytical, Worcestershire, UK) with 40 injections at 25°C. The experimental data were analyzed using MicroCal PEAQ-ITC analysis software with a single-site/double-site binding model.

### Phylogenetic analysis

We downloaded six predicted structures representing each type of GNAT acetyltransferase from the AlphaFold Protein Structure Database^53^ (AFDB) May 2024 release: P76594, A0A0M8Z1A2, O05581, D9SZU3, Q1D7F7 and P0A944 for type I, type II, cAMP-binding type III, ACT type III, NADP^+^-binding type III and type IV, respectively. Each structure was searched against AFDB50 (AFDB clustered by 50% 3Di sequence identity) using Foldseek^71^(version a2f62e) to obtain alignments with at least 90% of both 3Di sequences are covered (foldseek search -c 0.9 --cluster-search 1 --max-seqs 100000). Resulting alignments were then converted into Kraken2 sample report format and visualized as Sankey diagrams with Pavian^72^(version 1.0).

We collected whole genome assemblies of 300 bacterial species from NCBI GenBank 2024 update^73^, including *Escherichia coli*, *Streptomyces lividans*, *Micromonospora aurantiaca*, *Mycobacterium tuberculosis*, *Myxococcus xanthus* and 295 species randomly sampled across the bacterial domain. Four archaeal species under genus *Methanococcus* were included as an outgroup. Maximum likelihood tree was then inferred from the concatenation of amino acid alignments of 81 bacterial core genes extracted from the assemblies with UBCG^74^ (version 2.0), applying JTT+F+I+G evolutionary model of IQ-TREE^75^(version 2.2.2.7).

Finally, we’ve put the representative acetyltransferases predicted/experimental structures which has different types of regulatory domains: *Streptomyces lividans* PatA (*sl*PatA, AIJ12884.1) for Type II and *Escherichia coli* YiaC (*ec*YiaC, UniProt: P37664), while Type III includes *Mycobacterium tuberculosis* Pat (MtPat, PDB ID: 4AVB), *Micromonospora aurantiaca* PatB (*ma*PatB, UniProt: D9SZU3) and *Myxococcus xanthus* ActC (*mx*ActC, UniProt: Q1D7F7). *sl*PatA displays similarities to those of Type I with ligands, but the three of Type III GNATs bind to highly specific ligands, including cAMP, Cys/Arg and NADP^+^. Upon cAMP and Cys/Arg, *mt*Pat and *ma*PatB efficiently acetylate their substrates, respectively. But interestingly, *mx*ActC is inhibited when NADP^+^ is bound.

## DATA, MATERIALS, AND SOFTWARE AVAILABILITY

Atomic coordinates of the apo PatZ, liganded PatZ and ATP bound AGD of PatZ models were deposited in the RCSB PDB under accession numbers 9SIQ, 9IT0 and 9ISB, respectively. The cryoEM maps were deposited in the EMDB under accession numbers EMD-60849 (apo PatZ) and EMD-60853 (liganded PatZ). All code used for cryoEM data analysis, structure determination and refinement are publicly available.

## ACKNOWLEDGEMENTS

This work has been supported by the Korean National Research Foundation (2019R1C1C1004598, 2020R1A5A1018081, 2021M3A9I4021220, 2019M3E5D6063871, 2020R1A6C101A183) and SUHF foundation to S.-H.R and NRF-2018R1A5A1025077, NRF-2019R1A2C2004143, RS-2024-00335765 to Y.-J. S. The cryoEM data was collected and processed at The Center for Macromolecular and Cell Imaging (CMCI), Institute for Basic Science (IBS), Global Science experimental Data hub Center (GSDC) at Korea Institute of Science facilities. We thank members of CMCI for their discussions and suggestions. We thank members of the Frydman lab for useful discussions and suggestions. We thank the staff scientists for assistance at the beamline 5C of Pohang Light Source.

## AUTHOR INFORMATION

*School of Biological Sciences, Institute of Molecular Biology and Genetics, Seoul National University, Seoul, 08826, Republic of Korea*

Jun Bae Park, Yu-Yeon Han & Soung-Hun Roh

*School of Biological Sciences and Institute of Microbiology, Seoul National University, Seoul, 08826, Republic of Korea*

Gwanwoo Lee, Kyoo Heo & Yeong-Jae Seok

*Interdisciplinary Program in Bioinformatics, Seoul National University, Seoul, 08826, Republic of Korea*

Dongwook Kim

*Interdisciplinary Program in Bioinformatics, School of Biological Sciences, Institute of Molecular Biology and Genetics, Artificial Intelligence Institute, Seoul National University, Seoul, 08826, Republic of Korea*

Martin Steinegger

*Department of Systems Biology, College of Life Science and Biotechnology, Yonsei University, 50 Yonsei-ro, Seoul, 03722, Republic of Korea*

Hyun-Soo Cho

*Department of Chemistry, Chonnam National University, Gwangju, 61186, Republic of Korea*

Hyosuk Yun, Chul Won Lee

*School of Biological Sciences and Institute of Microbiology, Center for RNA Research, Institute for Basic Science, Seoul 08826, Republic of Korea*

Jeesoo Kim, Juhee Park & Jong-Seo Kim

## Author Contributions

S.-H.R. and Y.-J.S. conceived the project. J.B.P. and G.L. lead the project. J.B.P. and G.L. designed the experiment. J.B.P., G.L., Y.-Y.H. and K.H. performed molecular cloning, protein expression and purification of the proteins. CryoEM sample preparation, data collection and data processing were carried out by S.-H.R., J.B.P. and Y.-Y.H.. S.-H.R., J.B.P. and Y.-Y.H. analyzed the data, built and refined the model. For the X-ray crystallography, J.B.P. performed molecular cloning, protein expression, protein purification, crystallization, X-ray diffraction, data processing, model building and refinement under guidance of H.-S.C.. Y.-J.S. and G.L. conducted the enzymatic activity assay and biochemical assay. M.S. and D.K. performed phylogenetic analysis. All authors analyzed and discussed the results. J.B.P., G.L., Y.-Y.H. and D.K. wrote the paper with help from all authors.

## EXTENDED DATA and LEGENDS

**Extended Data Figure 1.**
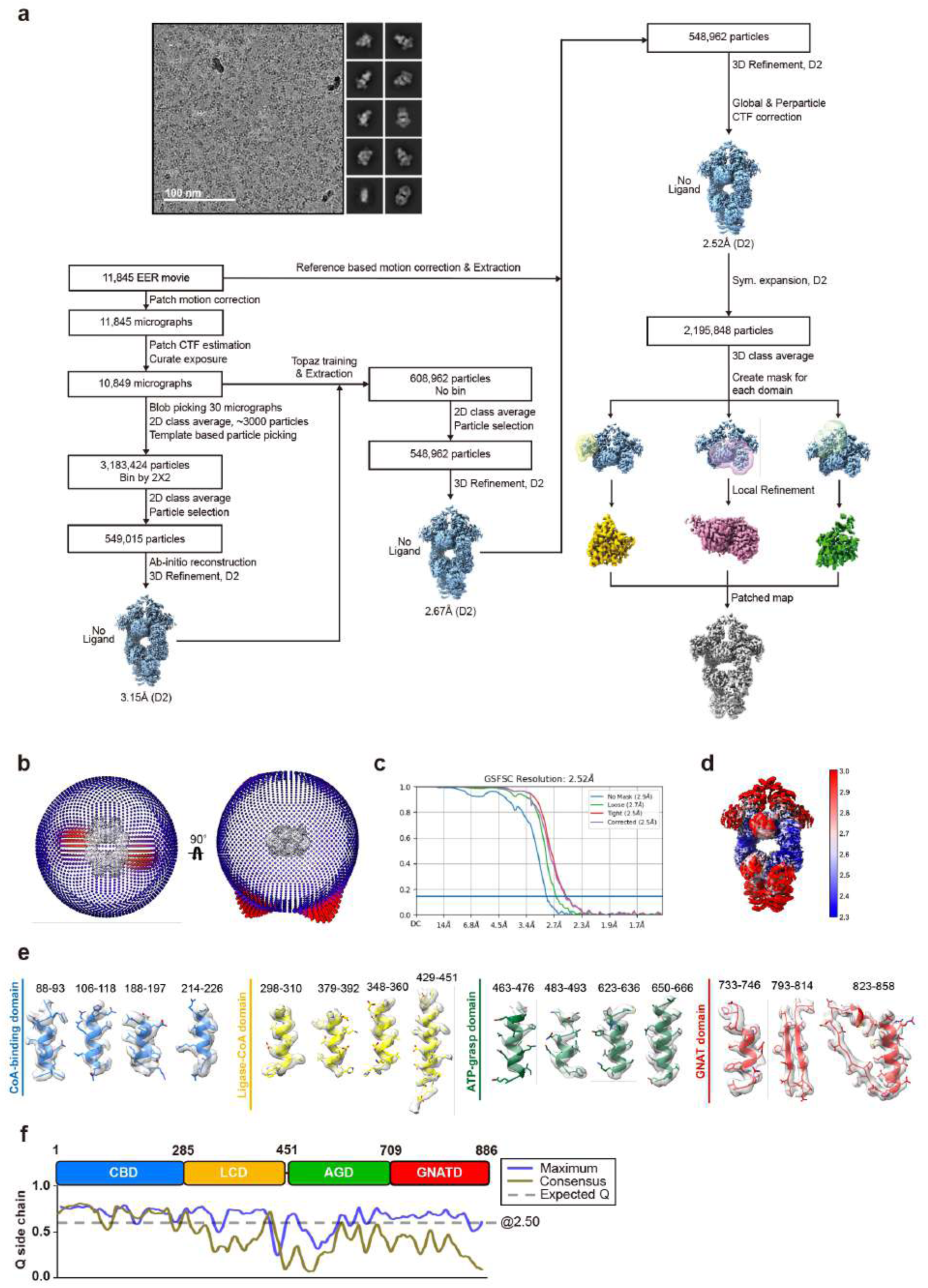
CryoEM processing workflow of apo-state PatZ. **a,** Schematic of data processing pipeline with representative cryogenic electron microscopy micrographs and two-dimensional averages. **b,** Angular distribution of views in the 3D reconstruction (left view and right view, rotated by 90°). **c,** Gold-standard fourier shell correlation (GSFSC) curve. **d,** Map in front view, colored according to local resolution. **e,** CryoEM density map features corresponding to the four subdomains. **f,** Side chain Q-score distribution across the four domains.

**Extended Data Figure 2.**
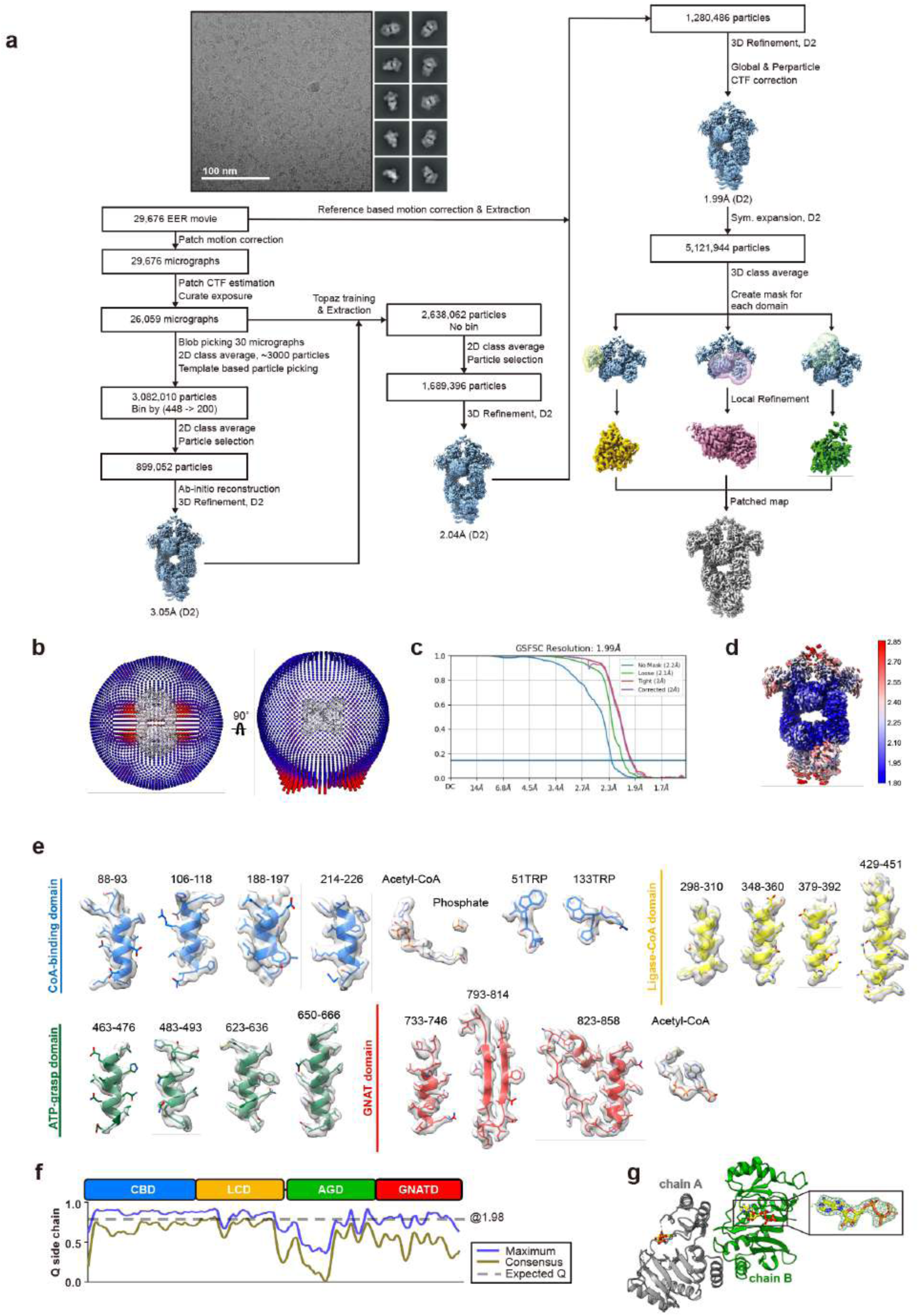
CryoEM processing workflow of liganded-PatZ and 2Fo-Fc electron density map of ATP in ATP bound AGD crystal structure. **a,** Schematic of data processing pipeline with representative cryogenic electron microscopy micrographs and two-dimensional averages. **b,** Angular distribution of views in the 3D reconstruction (left view and right view, rotated by 90°). **c,** Gold-standard fourier shell correlation (GSFSC) curve. **d,** Map in front view, colored according to local resolution. **e,** CryoEM density map features corresponding to the four subdomains. **f,** Side chain Q-score distribution across the four subdomains. **g,** Model refinement results for two AGD chains located in a single asymmetric unit of the crystal, with 2Fo-Fc electron density map of ATP contoured at 1σ.

**Extended Data Figure 3.**
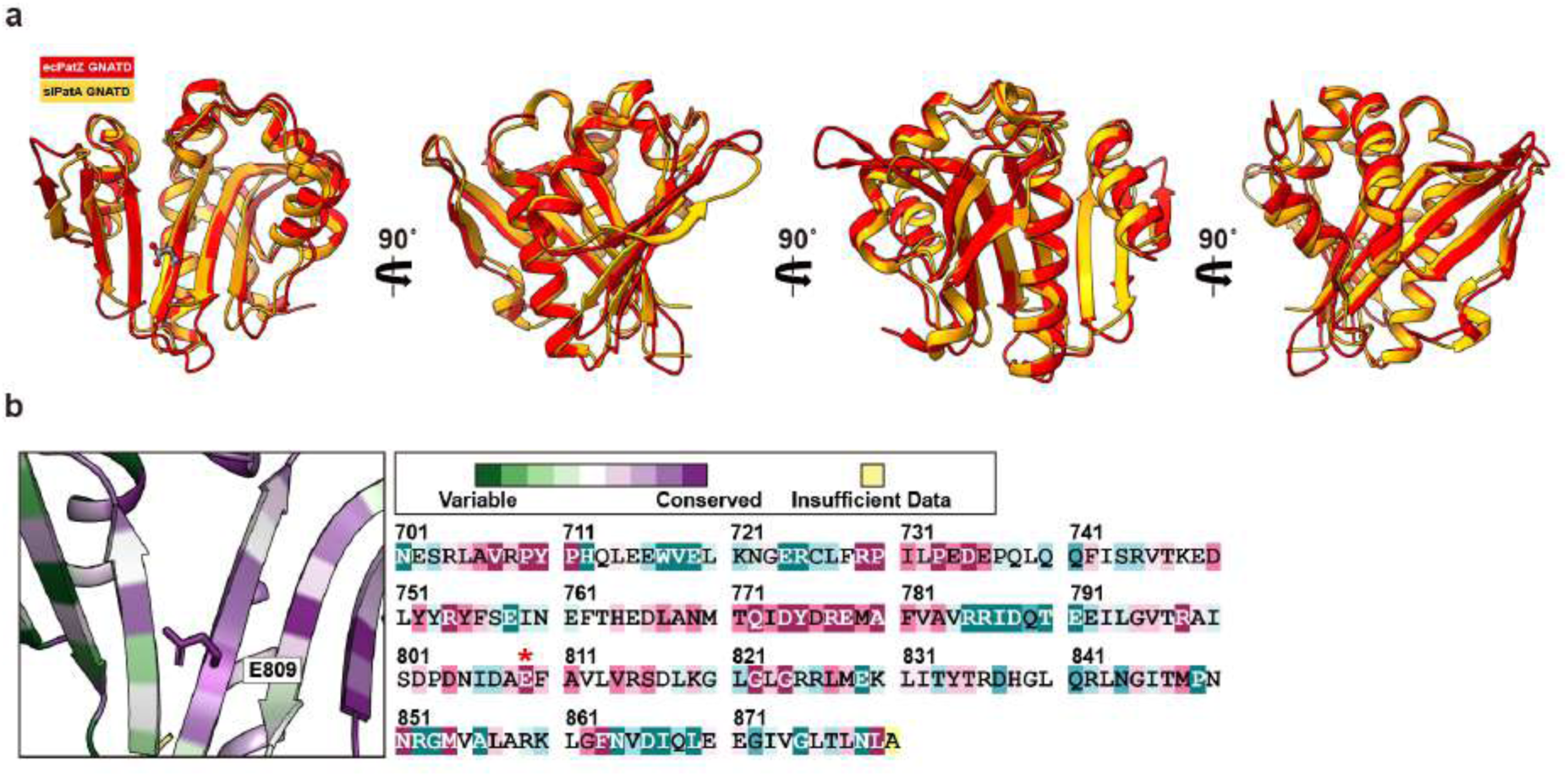
Structural similarity between GNATD of *ec*PatZ and *sl*PatA, and sequence conservation level of PatZ E809. **a,** Structural similarities between *ec*PatZ GNATD and *sl*PatA GNATD. **b,** ConSurf analysis of sequence conservation for PatZ E809.

**Extended Data Figure 4.**
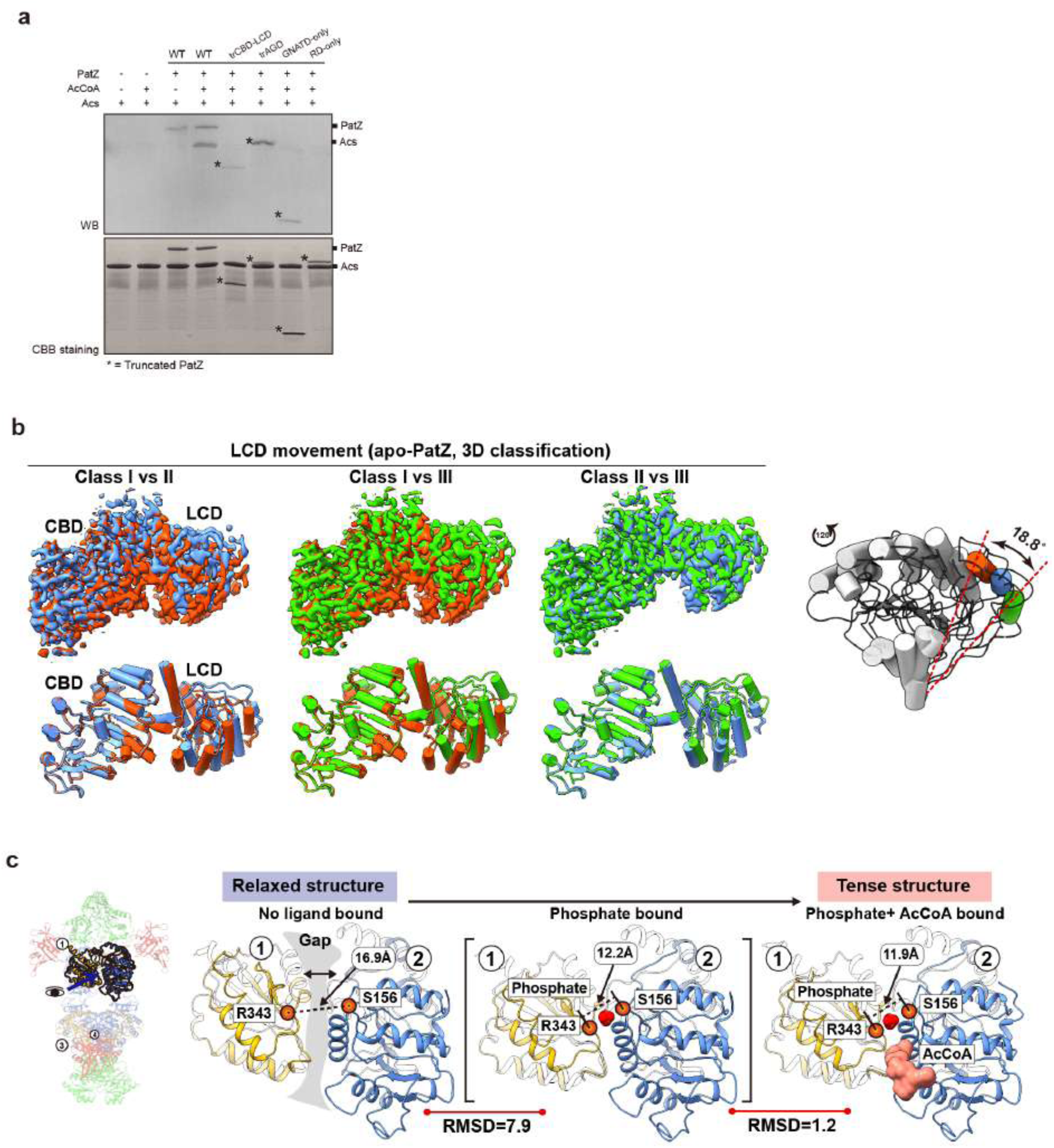
**a** Comparison of relative functional level between wild-type and truncated mutants. Asterisk mark refers to the truncated PatZs. **b,** Structural comparison of three distinct conformational states (Classes I-III) of apo-state PatZ LCD identified through 3D classification. **c,** Quantitative comparison of conformational changes in the regulatory domain induced by ligand binding.

**Extended Data Figure 5.**
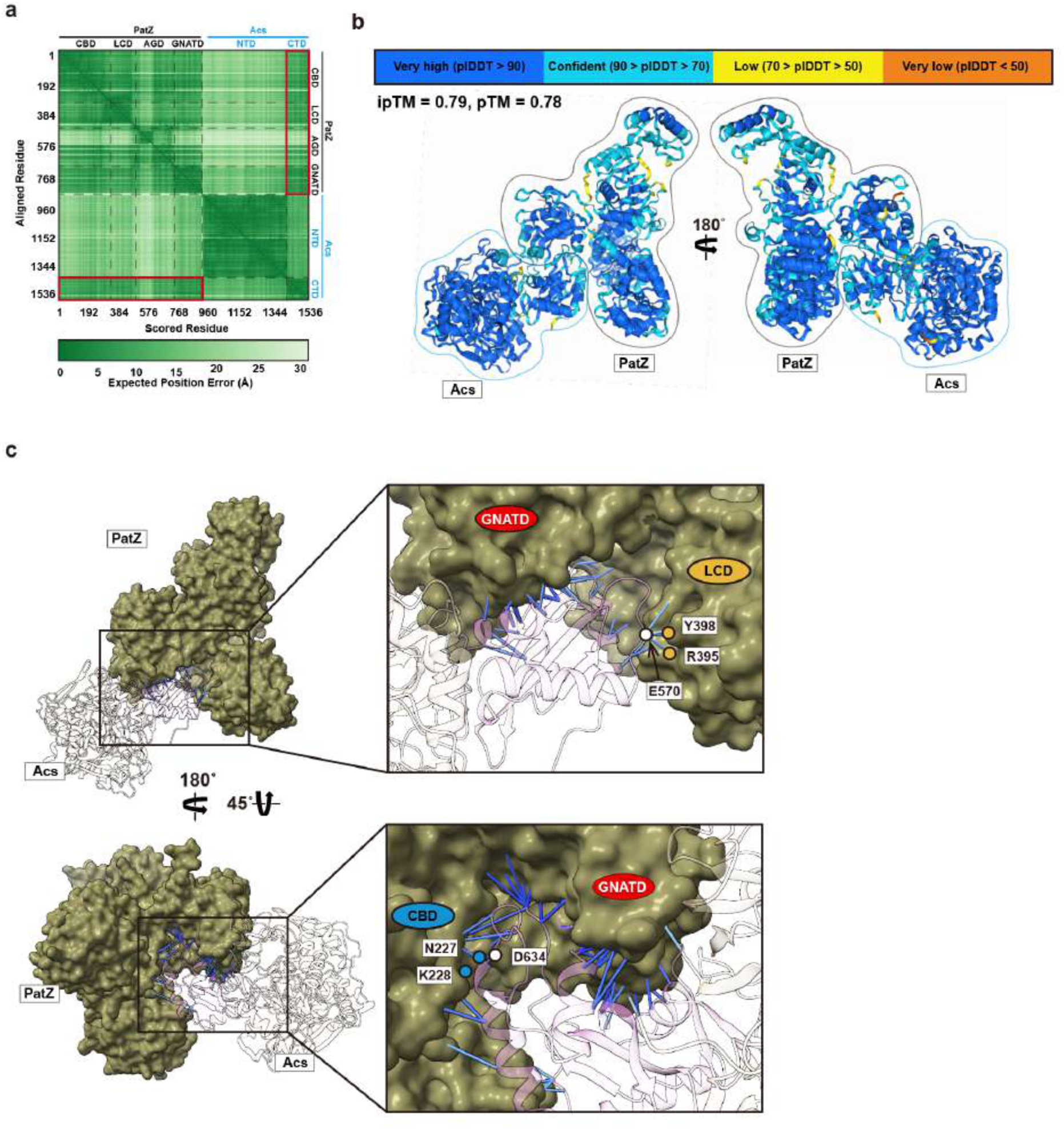
Prediction of PatZ-Acs complex structure using AlphaFold3. **a,** The Predicted Aligned Error (PAE) matrices calculated via AlphaFold3. **b,** pLDDT for all residues belonging to the Acs and PatZ. **c,** 3D visualization of prediction error estimation using PAE plot focused on PatZ-Acs interaction part. Stick style refers to residues originating from two chains within a 4 Å distance. The coloration follows the pLDDT color spectrum.

**Extended Data Figure 6.**
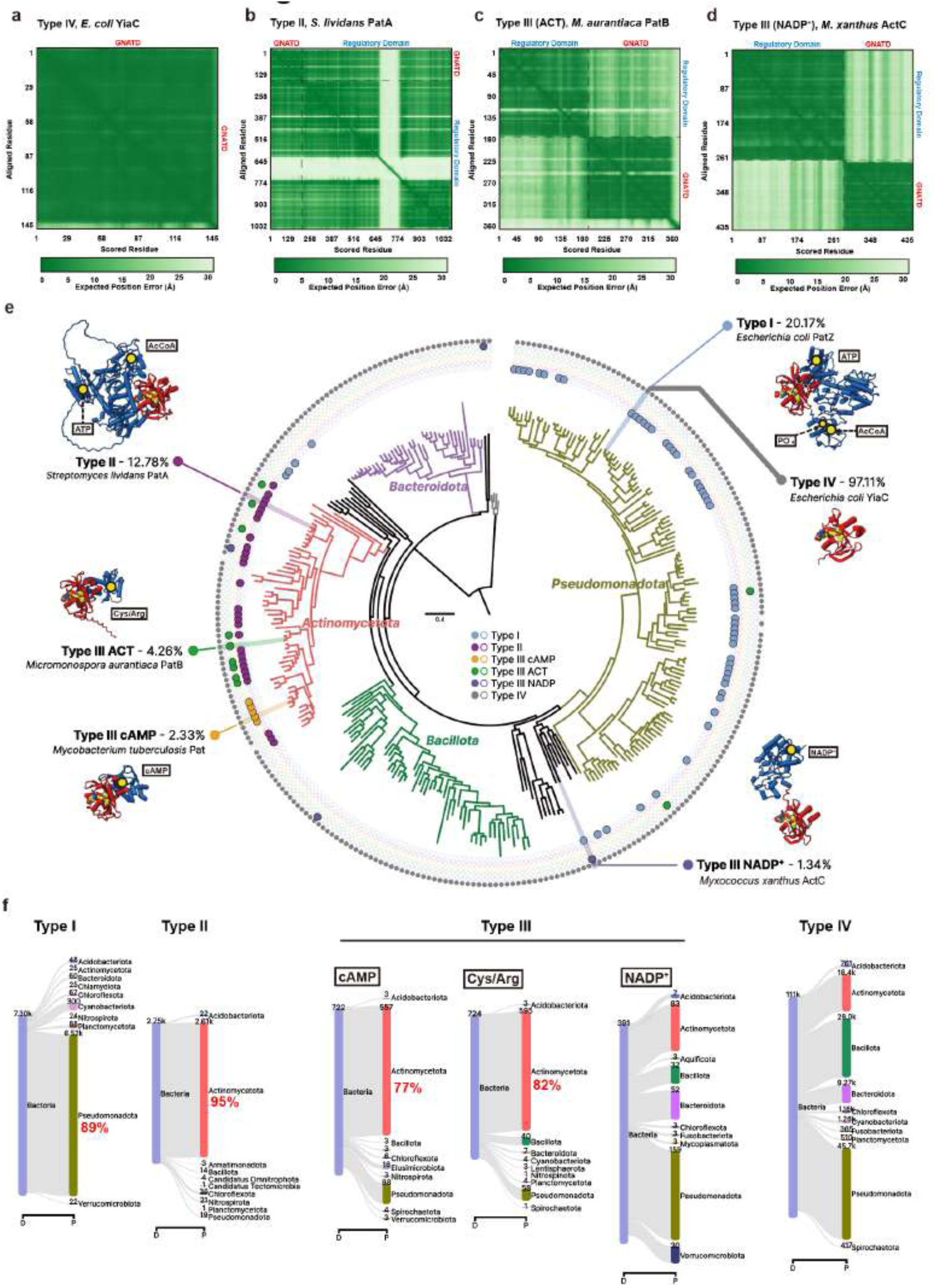
The PAE matrices that are calculated via AlphaFold3. **a,** *E.coli* YiaC (Type IV). **b,** *S. lividans* PatA (Type II). **c,** *M. aurantiaca* PatB (Type III). **d,** *M. xanthus* ActC (Type III). **e,** Domain-wide distribution of GNAT acetyltransferase types across a phylogenetic tree of 300 bacterial species. Different types are indicated by dots in the concentric circles, where a filled dot indicates the species containing a structure similar to the representative structure of each type. For each type, percentage of the species abundance among the bacterial domain as well as the representative structure are displayed on the location of corresponding species. **f**, Distribution of GNAT types across the bacterial domain displayed as Sankey plots. Shown are the ranks of domain (D) and phylum (P), with the top ten most abundant groups per rank; numbers indicate the amount of underlying species.

**Extended Data Figure 7.**
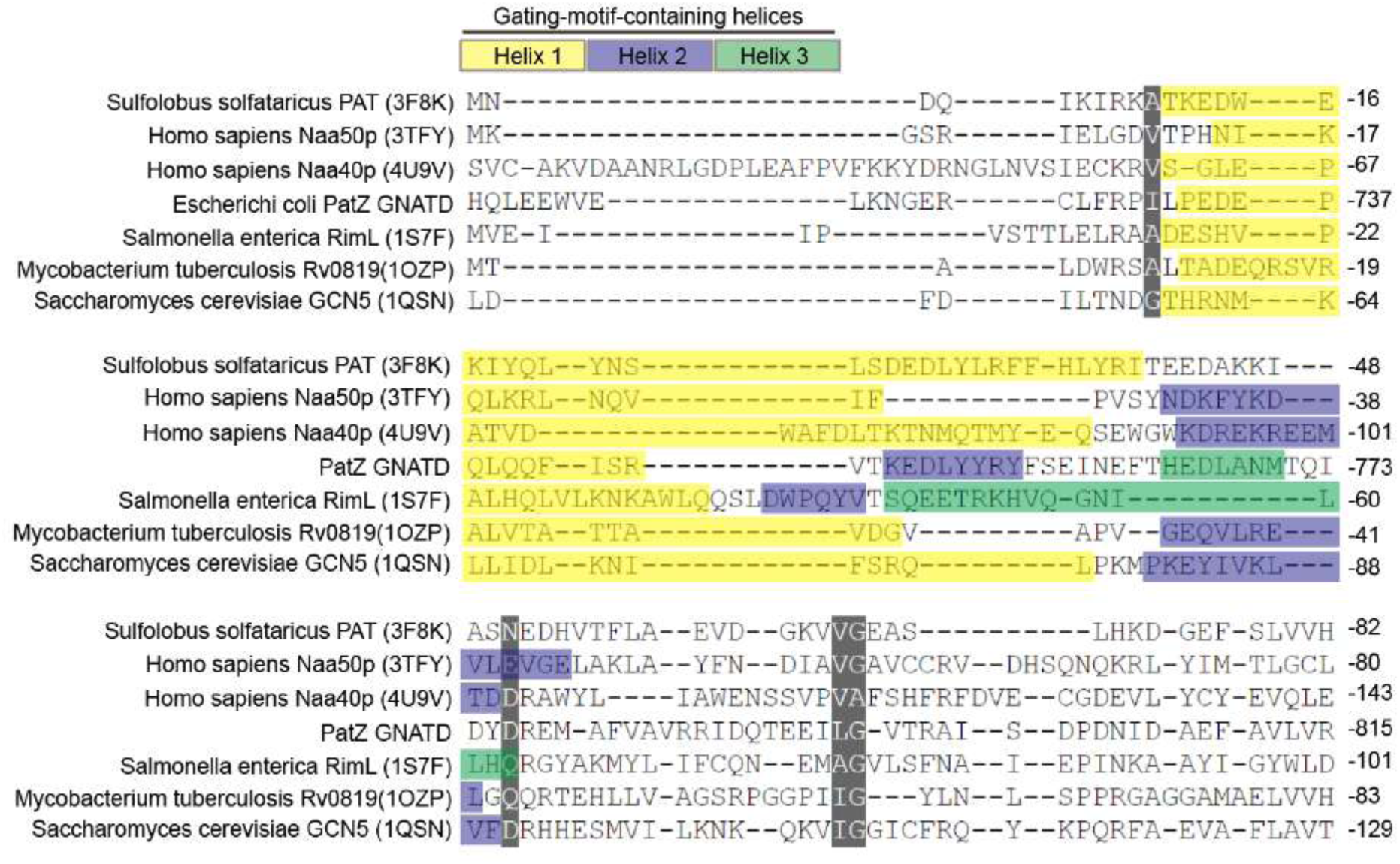
Structural similarity of gating-motif-containing helices. Comparative analysis of sequence-structure similarity in GNAT gating motifs across eukaryotes and prokaryotes. Distinct helices containing the gating motif are differentiated by color. Similarities among the sequences are given a gray background.

**Extended Data Figure 8.**
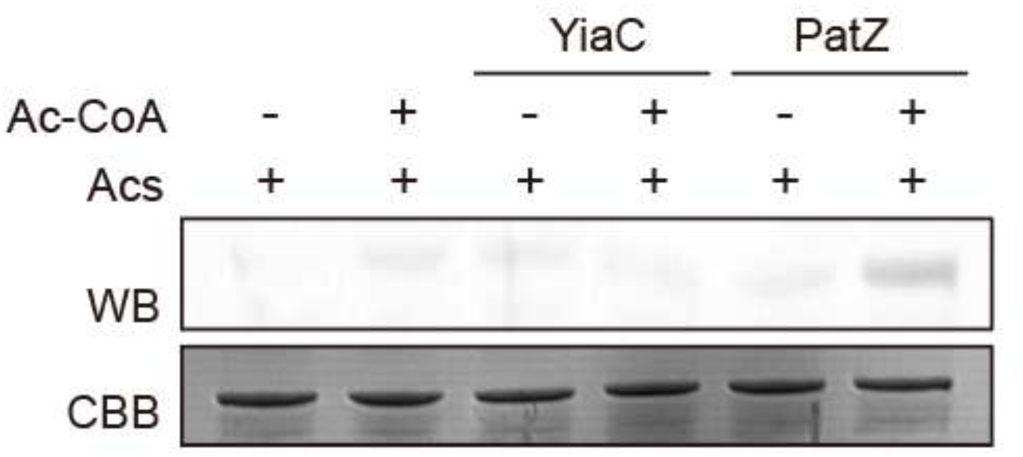
Western blot analysis using anti-acetylated lysine antibody for comparison of acetylation by type I *E. coli* PatZ and type IV *E. coli* YiaC.

**Supplementary Table 1.**
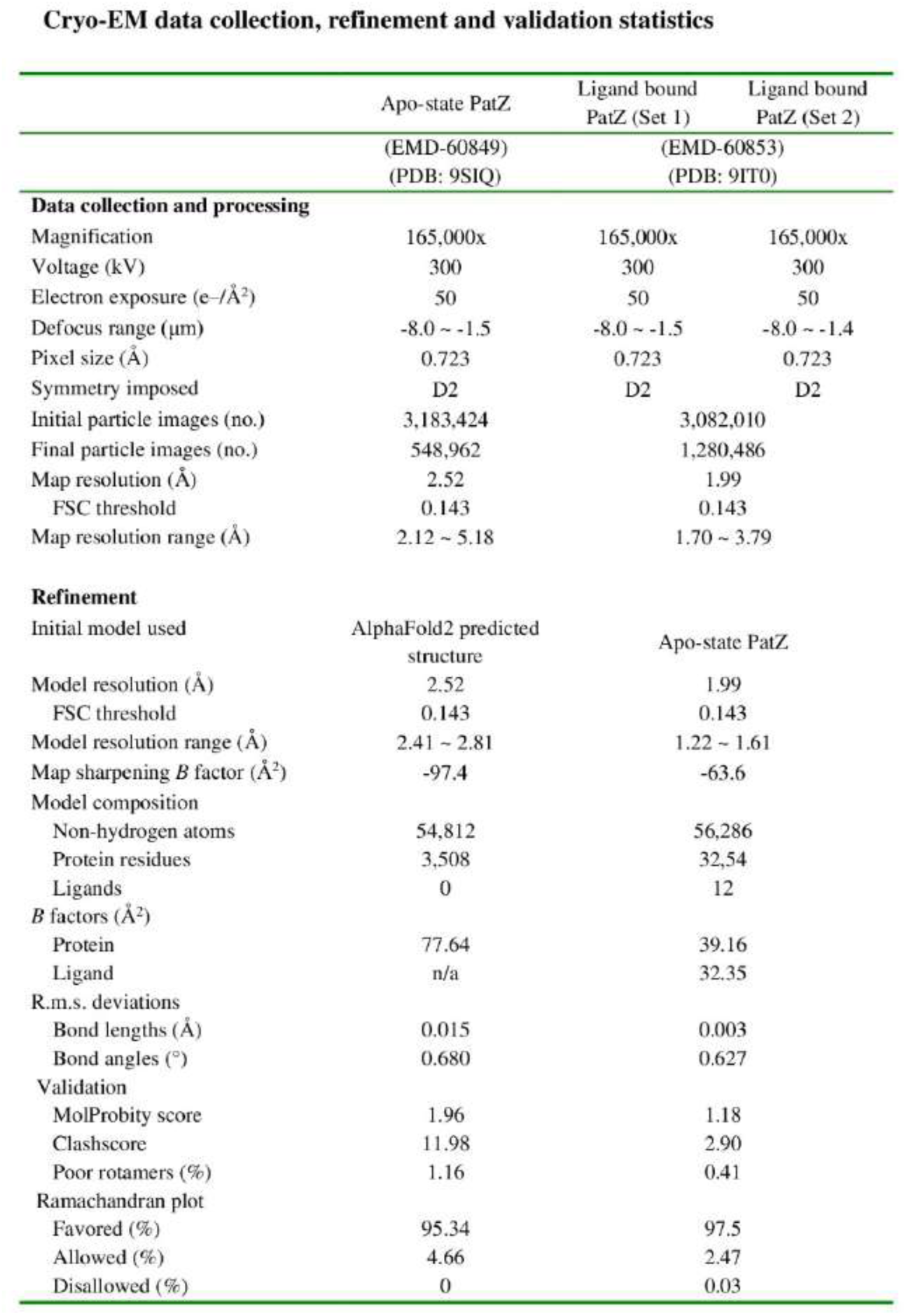
CryoEM data collection, refinement and validation statistics.

**Supplementary Table 2.**
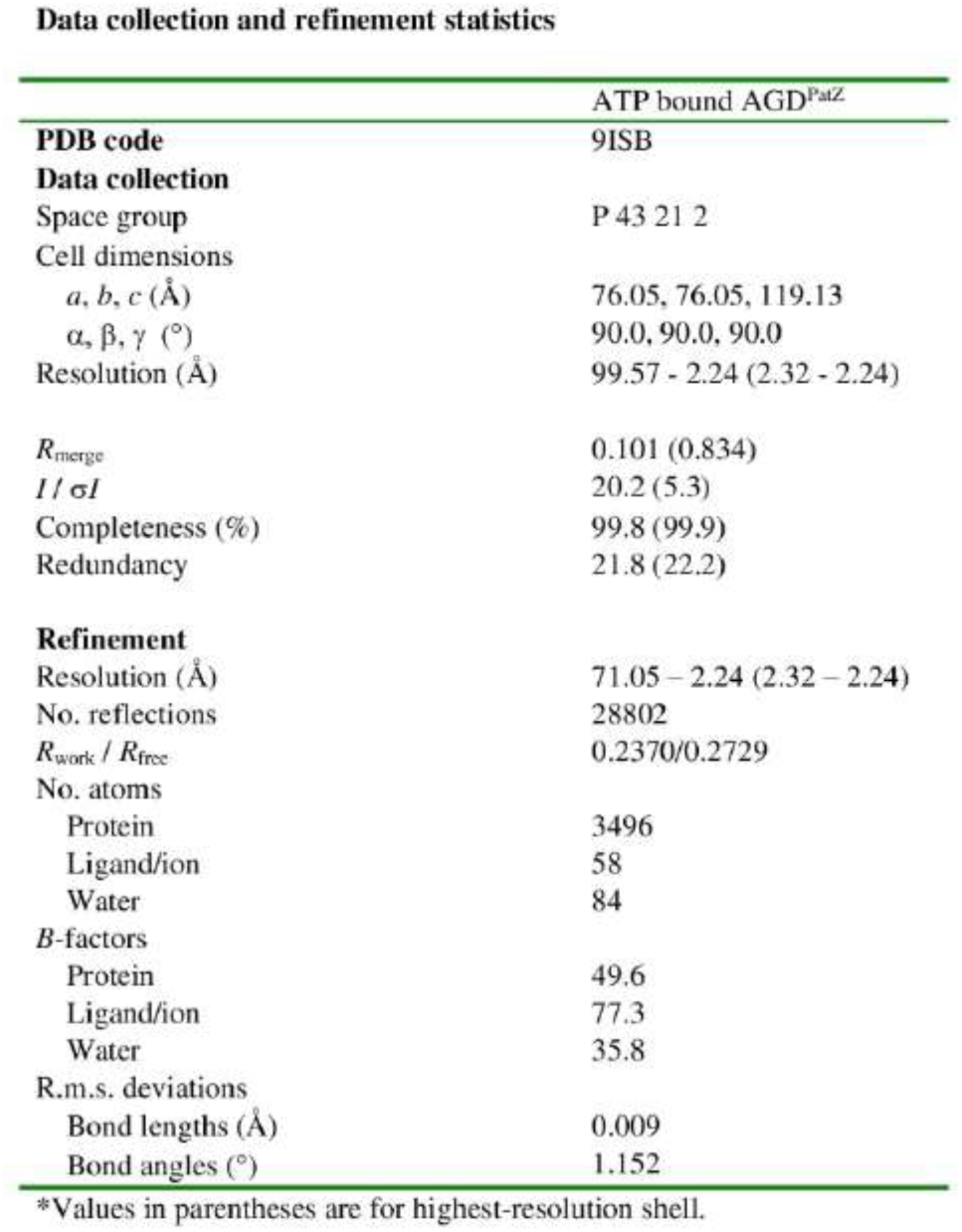
X-ray crystallographic data collection and refinement statistics.

**Supplementary video 1.** Tetramer assembly and subdomain composition of PatZ

**Supplementary video 2.** Ligands that bind to PatZ and their binding sites

**Supplementary video 3.** Ligand-mediated conformational change of PatZ GNAT domain

**Supplementary video 4.** Ligand-mediated global conformational change of PatZ

