## Supplementary data figure and tables for "Structural basis of the catalytic and allosteric mechanism of bacterial acetyltransferase PatZ"

### Extended Data Figure 1

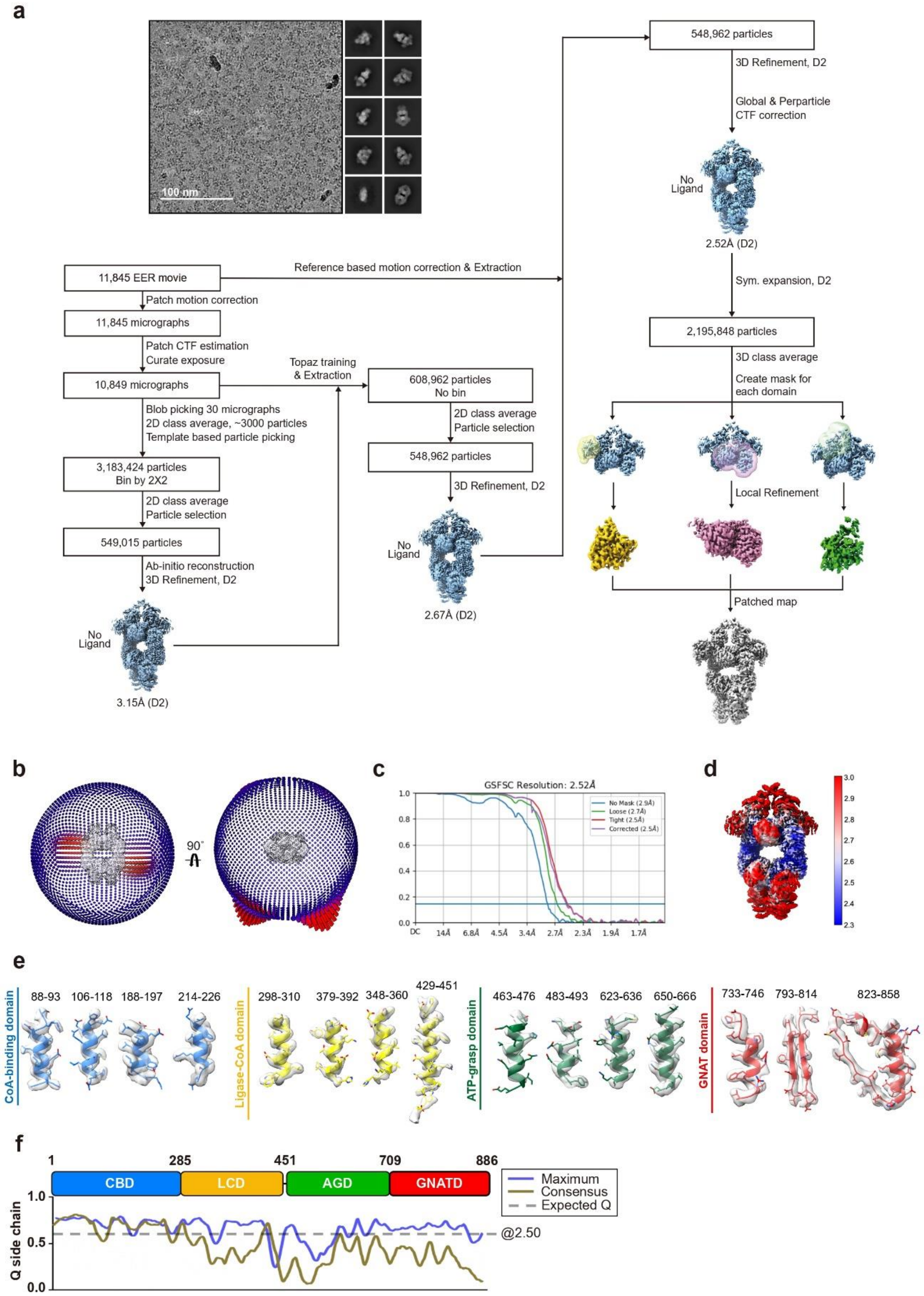

**Extended Data Figure 1.** CryoEM processing workflow of apo-state PatZ. **a,** Schematic of data processing pipeline with representative cryogenic electron microscopy micrographs and two-dimensional averages. **b,** Angular distribution of views in the 3D reconstruction (left view and right view, rotated by 90˚). **c,** Gold-standard fourier shell correlation (GSFSC) curve. **d,** Map in front view, colored according to local resolution. **e,** CryoEM density map features corresponding to the four subdomains. **f,** Side chain Q-score distribution across the four domains.

### Extended Data Figure 2

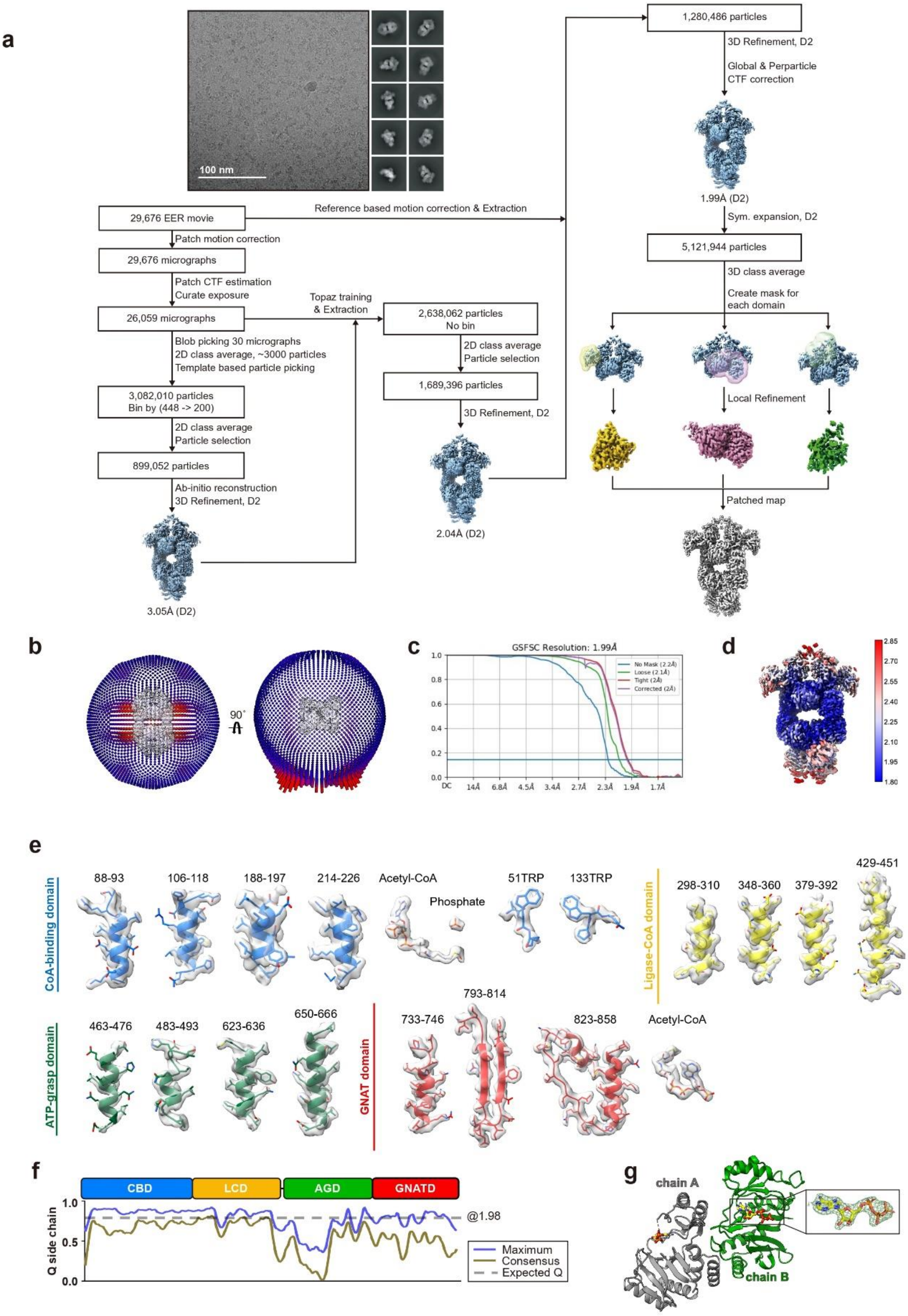

**Extended Data Figure 2.** CryoEM processing workflow of liganded-PatZ and 2Fo-Fc electron density map of ATP in ATP bound AGD crystal structure. **a,** Schematic of data processing pipeline with representative cryogenic electron microscopy micrographs and two-dimensional averages. **b,** Angular distribution of views in the 3D reconstruction (left view and right view, rotated by 90˚). **c,** Gold-standard fourier shell correlation (GSFSC) curve. **d,** Map in front view, colored according to local resolution. **e,** CryoEM density map features corresponding to the four subdomains. **f,** Side chain Q-score distribution across the four subdomains. **g,** Model refinement results for two AGD chains located in a single asymmetric unit of the crystal, with 2Fo-Fc electron density map of ATP contoured at 1σ.

#### Extended Data Figure 3

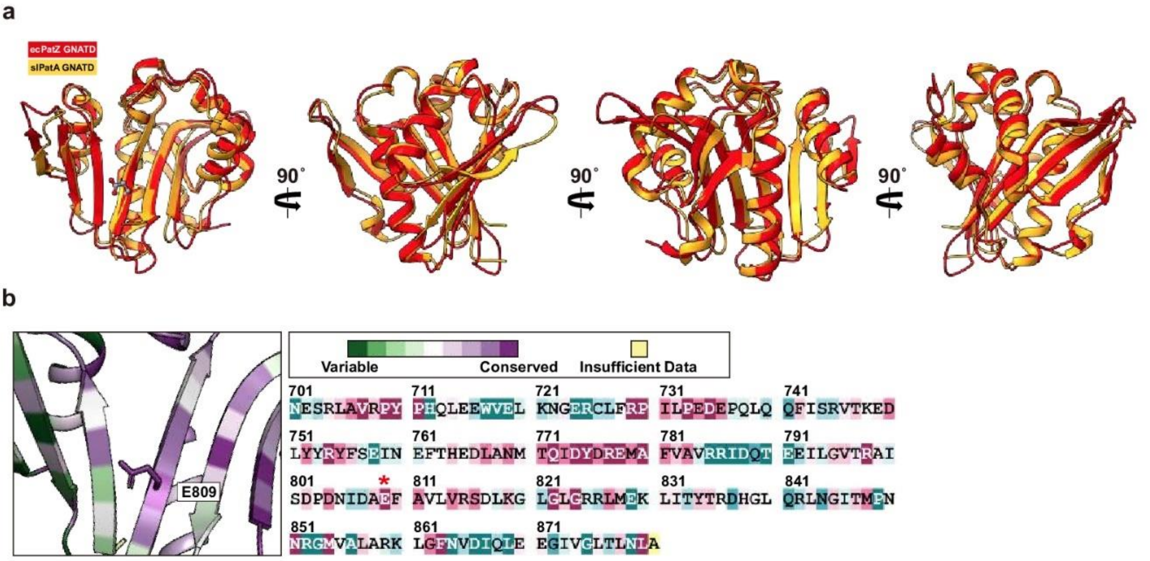


**Extended Data Figure 3.** Structural similarity between GNATD of *ec*PatZ and *sl*PatA, and sequence conservation level of PatZ E809. **a,** Structural similarities between *ec*PatZ GNATD and *sl*PatA GNATD. **b,** ConSurf analysis of sequence conservation for PatZ E809.

### Extended Data Figure 4

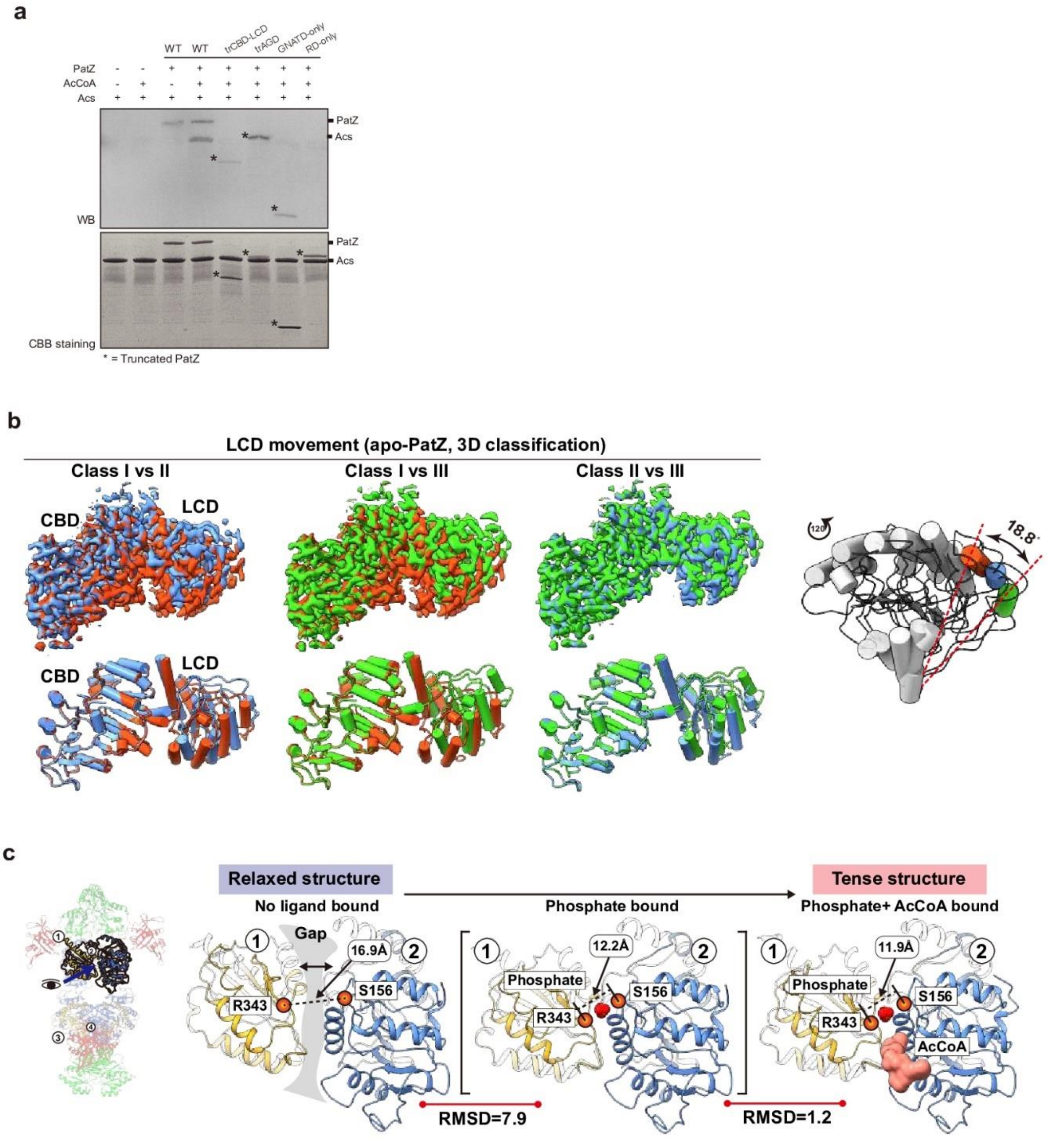


**Extended Data Figure 4.** **a** Comparison of relative functional level between wild-type and truncated mutants. Asterisk mark refers to the truncated PatZs. **b,** Structural comparison of three distinct conformational states (Classes I-III) of apo-state PatZ LCD identified through 3D classification. **c,** Quantitative comparison of conformational changes in the regulatory domain induced by ligand binding.

### Extended Data Figure 5

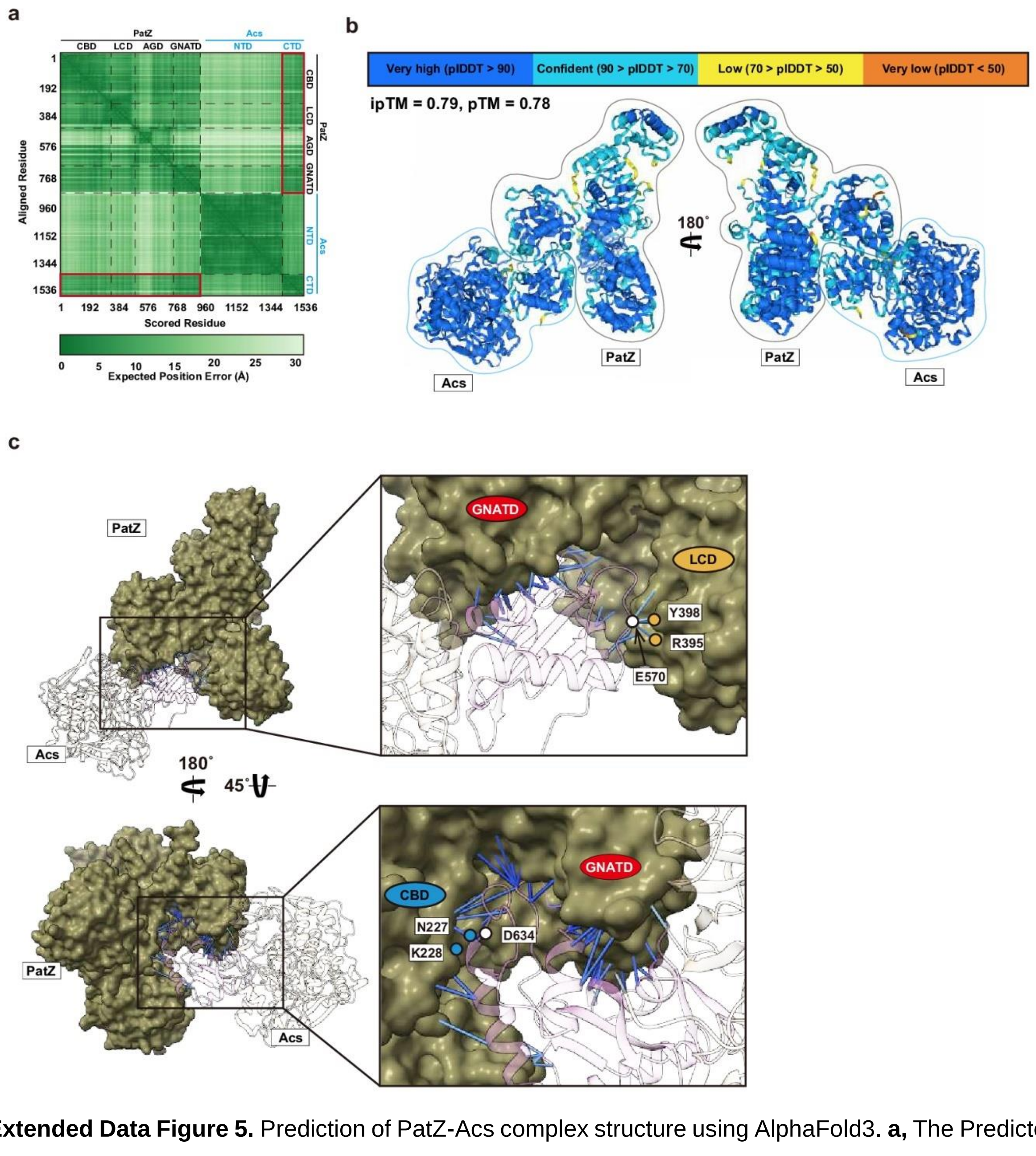


**Extended Data Figure 5.** Prediction of PatZ-Acs complex structure using AlphaFold3. **a,** The Predicted Aligned Error (PAE) matrices calculated via AlphaFold3. **b,** pLDDT for all residues belonging to the Acs and PatZ. **c,** 3D visualization of prediction error estimation using PAE plot focused on PatZ-Acs interaction part. Stick style refers to residues originating from two chains within a 4 Å distance. The coloration follows the pLDDT color spectrum.

#### Extended Data Figure 6

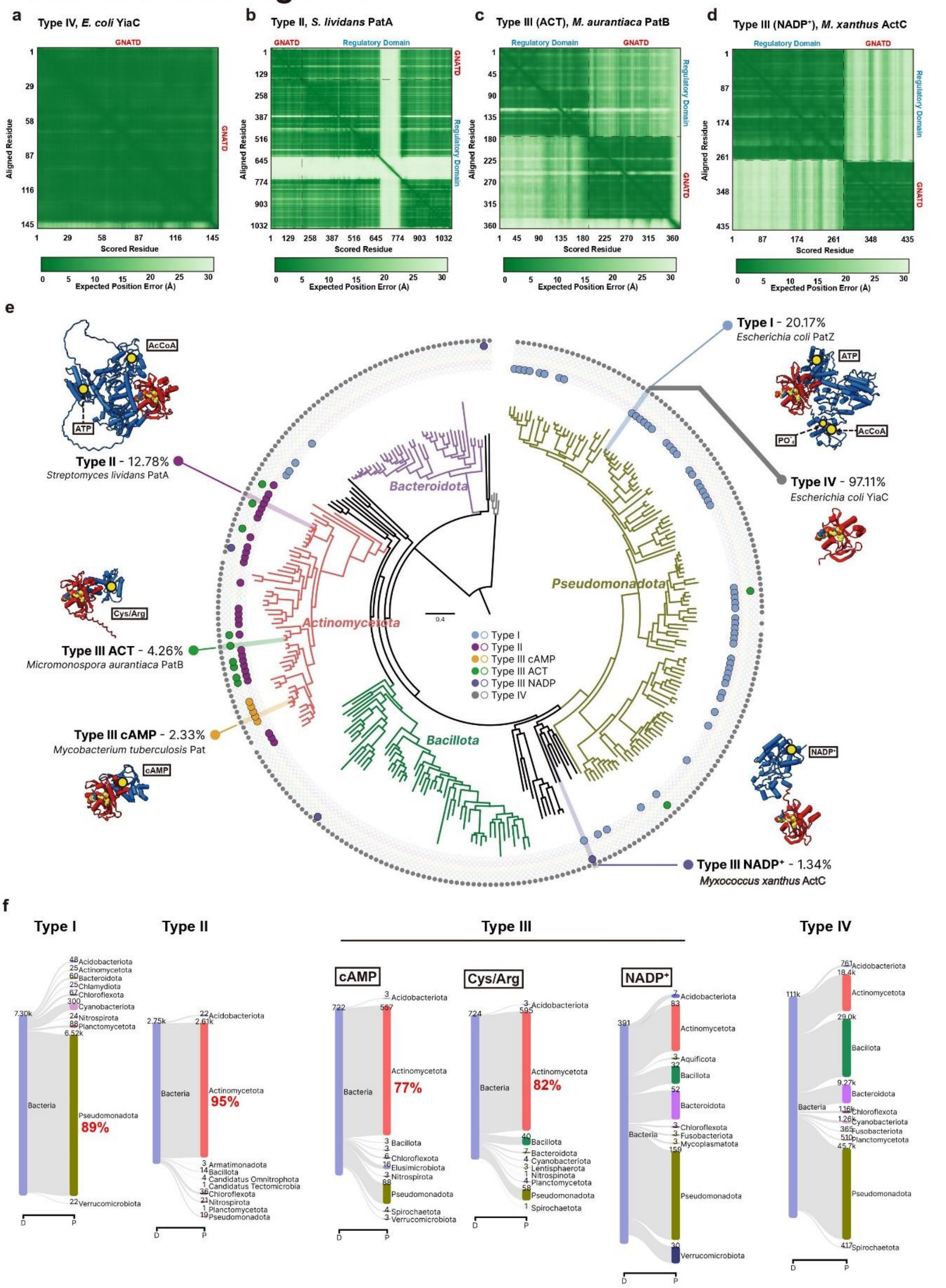


**Extended Data Figure 6.** The PAE matrices that are calculated via AlphaFold3. **a,** *E.coli* YiaC (Type IV). **b,** *S. lividans* PatA (Type II). **c,** *M. aurantiaca* PatB (Type III). **d,** *M. xanthus* ActC (Type III). **e,**

Domain-wide distribution of GNAT acetyltransferase types across a phylogenetic tree of 300 bacterial species. Different types are indicated by dots in the concentric circles, where a filled dot indicates the species containing a structure similar to the representative structure of each type. For each type, percentage of the species abundance among the bacterial domain as well as the representative structure are displayed on the location of corresponding species. **f**, Distribution of GNAT types across the bacterial domain displayed as Sankey plots. Shown are the ranks of domain (D) and phylum (P), with the top ten most abundant groups per rank; numbers indicate the amount of underlying species.

### Extended Data Figure 7

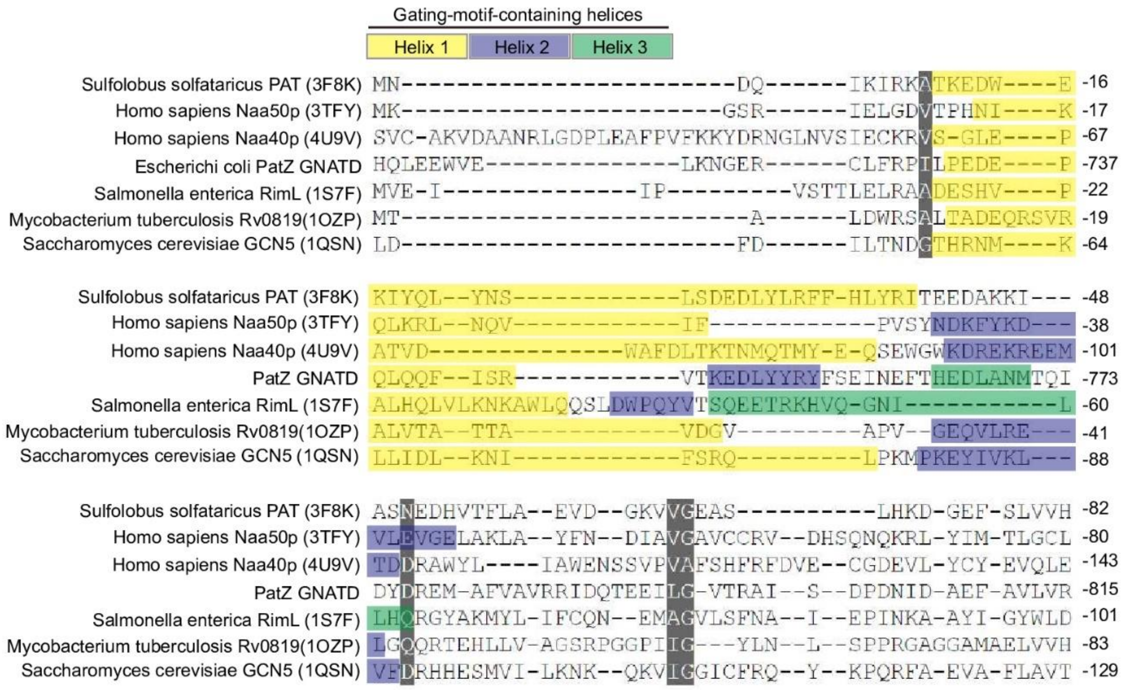


**Extended Data Figure 7.** Structural similarity of gating-motif-containing helices. Comparative analysis of sequence-structure similarity in GNAT gating motifs across eukaryotes and prokaryotes. Distinct helices containing the gating motif are differentiated by color. Similarities among the sequences are given a gray background.

#### Extended Data Figure 8

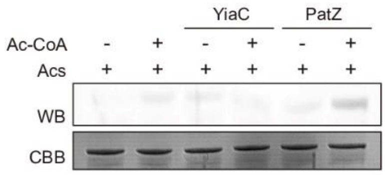


**Extended Data Figure 8**. Western blot analysis using anti-acetylated lysine antibody for comparison of acetylation by type I *E. coli* PatZ and type IV *E. coli* YiaC.

**Supplementary Table 1.** CryoEM data collection, refinement and validation statistics

**Cryo-EM data collection, refinement and validation statistics**

| | Apo-state PatZ | Ligand bound PatZ (Set 1) | Ligand bound PatZ (Set 2) |
|---|---|---|---|
| | (EMD-60849) (PDB: 9SIQ) | (EMD-60853) (PDB: 9IT0) | |
| **Data collection and processing** | | | |
| Magnification | 165,000x | 165,000x | 165,000x |
| Voltage (kV) | 300 | 300 | 300 |
| Electron exposure (e–/Å$^2$) | 50 | 50 | 50 |
| Defocus range (μm) | -8.0 ~ -1.5 | -8.0 ~ -1.5 | -8.0 ~ -1.4 |
| Pixel size (Å) | 0.723 | 0.723 | 0.723 |
| Symmetry imposed | D2 | D2 | D2 |
| Initial particle images (no.) | 3,183,424 | 3,082,010 | |
| Final particle images (no.) | 548,962 | 1,280,486 | |
| Map resolution (Å) | 2.52 | 1.99 | |
| FSC threshold | 0.143 | 0.143 | |
| Map resolution range (Å) | 2.12 ~ 5.18 | 1.70 ~ 3.79 | |
| **Refinement** | | | |
| Initial model used | AlphaFold2 predicted structure | Apo-state PatZ | |
| Model resolution (Å) | 2.52 | 1.99 | |
| FSC threshold | 0.143 | 0.143 | |
| Model resolution range (Å) | 2.41 ~ 2.81 | 1.22 ~ 1.61 | |
| Map sharpening *B* factor (Å$^2$) | -97.4 | -63.6 | |
| Model composition | | | |
| Non-hydrogen atoms | 54,812 | 56,286 | |
| Protein residues | 3,508 | 32,54 | |
| Ligands | 0 | 12 | |
| *B* factors (Å$^2$) | | | |
| Protein | 77.64 | 39.16 | |
| Ligand | n/a | 32.35 | |
| R.m.s. deviations | | | |
| Bond lengths (Å) | 0.015 | 0.003 | |
| Bond angles (°) | 0.680 | 0.627 | |
| Validation | | | |
| MolProbity score | 1.96 | 1.18 | |
| Clashscore | 11.98 | 2.90 | |
| Poor rotamers (%) | 1.16 | 0.41 | |
| Ramachandran plot | | | |
| Favored (%) | 95.34 | 97.5 | |
| Allowed (%) | 4.66 | 2.47 | |
| Disallowed (%) | 0 | 0.03 | |

**Supplementary Table 2.** X-ray crystallographic data collection and refinement statistics

**Data collection and refinement statistics**

| | ATP bound AGD$^{PatZ}$ |
|---|---|
| **PDB code** | 9ISB |
| **Data collection** | |
| Space group | P 43 21 2 |
| Cell dimensions | |
| $a$, $b$, $c$ (Å) | 76.05, 76.05, 119.13 |
| α, β, γ (°) | 90.0, 90.0, 90.0 |
| Resolution (Å) | 99.57 - 2.24 (2.32 - 2.24) |
| $R_{merge}$ | 0.101 (0.834) |
| $I / \sigma I$ | 20.2 (5.3) |
| Completeness (%) | 99.8 (99.9) |
| Redundancy | 21.8 (22.2) |
| **Refinement** | |
| Resolution (Å) | 71.05 – 2.24 (2.32 – 2.24) |
| No. reflections | 28802 |
| $R_{work}$ / $R_{free}$ | 0.2370/0.2729 |
| No. atoms | |
| Protein | 3496 |
| Ligand/ion | 58 |
| Water | 84 |
| $B$-factors | |
| Protein | 49.6 |
| Ligand/ion | 77.3 |
| Water | 35.8 |
| R.m.s. deviations | |
| Bond lengths (Å) | 0.009 |
| Bond angles (°) | 1.152 |

*Values in parentheses are for highest-resolution shell.
